# Bench-ML: A Benchmarking Web Interface for Machine Learning Methods and Models in Genomics

**DOI:** 10.1101/2023.06.05.543750

**Authors:** Lopez Rene, Makita Mario, Ortega Laura, Lal Avantika, De La Garza Hernan

**Affiliations:** Instituto Tecnologico de Chihuahua 2; Stanford University

**Keywords:** Machine Learning, Deep Learning, Model Evaluation, Benchmarking, Genomics

## Abstract

Machine learning is a complex but essential technology in genomics data analysis and its popularity has increased the rate of new methodological approaches published but this raises the question of how models should be benchmarked and validated.

Bench-ML is a generalizable and easy to use web interface for benchmarking and validation that can preprocess data, train, test, evaluate and compare machine learning algorithms for genomics. It makes benchmarking machine learning methods more accessible by enabling genomics scientists to perform end-to-end analyses, visualize results and evaluate performance or metrics to compare methods and models by providing a point of reference using only a web browser.

***To improve something it needs to be measured***; To benchmark and evaluate models Bench-ML provides several strategies, methodologies, and tools to generate measurements and visualizations to track experiments to help identify areas of opportunity using metrics such as loss and accuracy, model visualization, learning and saturation curves, principal component analysis, feature scoring, confusion matrix, regression for training and test data, mean absolute error, etc.

Bench-ML explains the different options to test and validate machine and deep learning models to identify problematic areas and potentially improve performance. Bench-ML provides several strategies to improve performance like showing when a model is not performing or when different hyperparameters values could be needed, it also helps fine tune hyperparameter values and to identify accuracy across multiple classes and from these classes which class could affect performance.

The selection, development, and comparison of machine learning methods and models in genomics datasets can be a daunting task based on the goals of a particular study or the target problem. Machine learning is very good at pattern recognition but modeling the world is much more than that so how to know if a machine learning method or model is performing at a good sensitivity and specificity in large genomics datasets is still a big problem and this is where Bench-ML can help.

## Introduction

Sequencing or genomics has quickly grown into a new approach for clinical diagnosis of disease. Next-generation sequencing technologies and clinical interpretation of the variants are used today to guide health care providers in understanding and diagnosing their patient’s disease (4).

The tremendous amount of biological sequence data available, combined with the recent methodological breakthrough in deep learning in domains such as computer vision or natural language processing, is leading today to the transformation of bioinformatics through the emergence of deep genomics, the application of deep learning to genomic sequences (10).

Machine learning (ML) is a field of study concerned with algorithms that learn from examples and is divided into supervised and unsupervised learning. This article mostly focuses on supervised machine learning given that in genomics there are many datasets to learn from. Supervised machine learning examples are given by labels and features, the labels in genomics are the classes machine learning is trying to predict or classify. Features in genomics could be human gene expression values as in RNA-Seq analysis, DNA variants as used in phenotype/genotype associations, species or strains expression values as used in microbiome 16s or metagenomics or metatranscriptomics studies, DNA sequences to identify downregulated or upregulated regions, etc.

There has been an incoming amount of work and papers with deep learning in new fields such as dentistry and biomedicine that has helped researchers enhance their research (15); but this highlights the need for new benchmarking tools to evaluate their studies and results better.

There have been a number of new genomics diagnostic tools developed using machine learning methods but irrespective of how a ML genomic tool is derived it must be properly evaluated. In (16) the authors describe how to evaluate tools but it doesn’t cover how to check if the resulting model is the best approach or if the model could be improved or if the model has any bias or if it performs with the same level of accuracy for all classes; these are the cases where benchmarking and comparing models and methods evaluations in detail could help identify areas that needs improvement.

There are also many tools being upgraded to deep learning from traditional statistical methods such as the “SNP and small-indel variant caller using deep neural networks” (2). And for these reasons Machine learning has become an essential tool in biomedicine to make sense of large, high-dimensional datasets such as those found in genomics, proteomics, and imaging datasets (32). Some libraries have been developed but without a web interface or not suitable for genomics applications (9).

While some machine learning algorithms can work with categorical data most algorithms don’t, for efficiency reasons. One of the aims of a benchmarking tool is to facilitate data preparation and categorical data often needs to be converted to either ordinal encoding, one-hot encoding or dummy variable encoding in order to use numerical data. For DNA data types which are composed of nucleotides Adenine (A), Thymine (T), Cytosine (C), Guanine (G) one-hot encoding is preferred to avoid an ordinal relationship where the number values could influence training the model.

In clinical genomics it is required that every sample is described in detail. What was the measurement, is it valid, why it didn’t perform, why did we get that result, would all the samples within that class or category perform at the same level of accuracy, etc. More than often we develop a model but we don’t know if our model converged or if it could be improved due to lack of knowledge or adequate evaluation tools. In some settings only the total percent of sensitivity is described like in a clinical cancer dataset where sensitivity of plasma-derived NGS was 85.0%, comparable to 80.7% sensitivity for tissue (5) but there is no mention if any classes or types of cancer underperformed given that there are 33 different types of cancer and is very unlikely that all of them performed at 85% sensitivity from the plasma-derived samples.

Many machine learning methods suppose that random observations in a problem are not dependent on each other and have a constant probability of occurring because it does large scale pattern recognition on suitable collected independent and identically distributed data (20). Although, for humans it is easy to make inferences about causal relations between different elements but until now machine learning has neglected a full integration of causality because there are many variables in a dataset. System benchmarking and visualizing our models could show if a model isn’t properly fit and if the data or causality could potentially play a role in creating bias. In “towards causal representation of machine learning” the authors argue that machine learning will benefit from integrating causal concepts (20).

Research on artificial neural networks was motivated by the observation that human intelligence emerges from highly parallel networks of relatively simple, non-linear neurons that learn by adjusting the strengths of their connections. Deep neural networks also known as Deep Learning (DL) uses many layers of activity vectors as representations and learning the connections strengths that give rise to these vectors by following the stochastic gradient of an objective function that measures how well the network is performing and the key is handling large training sets or “depth” such as those from genomics because shallow networks do not work so well (22).

The question is how can neural networks learn the rich internal representations required for difficult tasks such as understanding DNA, gene expression, or recognizing objects and how can we identify those. This is why Bench-ML provides an easy to follow web interface with methods, models, parameters, with ways to execute those models, understand and visualize the results. In supervised machine learning, large high-dimensional datasets are used to build statistical models that can predict discrete classes or labels as a classification analysis or continuous values like in regression analysis (32). Example applications of ML to biomedicine include developing predictive models for fusion methods for cancer (1), drug metabolism rates (33), genotype-phenotype associations (34), blood metabolome predicting guts microbiome alfa-diversity (7), drug metabolic activity (6), radiotherapy targeting of non-small-cell lung cancer (3), epigenomics (42), and drug response in model systems (35). Deep learning, which leverages multi-layer neural networks, has been used for prediction of splice sites (36), protein structures (37), and cancer diagnosis from histopathology images (38).

Deep learning (DL) is a form of machine learning that does not require explicit hard coding. The architecture of deep neural networks is inspired by neuroscience and the biological brain. Like the biological brain, the inner workings of exactly why deep networks work are largely unexplained and there is no single unifying theory (18) which makes DL harder to validate and benchmark.

Neural networks typically consist of many layers connected via many nonlinear intertwined relations. Even if one is to inspect all these layers and describe their relations, it is unfeasibly to fully comprehend how the neural network came to its decision. Therefore, deep learning is often considered a ‘black box’. Concern is mounting in various fields of application that these black boxes may be biased in some way, and that such bias goes unnoticed. Especially in medical applications, this can have far-reaching consequences (11).

There has been a call for approaches to better understand the black box. Such approaches are commonly referred to as interpretable deep learning or explainable artificial intelligence (XAI). These terms are commonly interchanged (12)(13).

Genomics datasets have much more dimensions compared to other fields; and for certain types of compositional functions, it is proven that deep networks of the convolutional type (even without weight sharing) can avoid the curse of dimensionality (18).

Deep networks tend to be over-parameterized and perform well on out-of-sample data. It is demonstrated that for classification problems, given a standard deep network, trained with gradient descent algorithms, it is the direction in the parameter space that matters, rather than the norms or the size of the weights (18). In general, most of the existing convolutional neural network (CNN)-based deep-learning models suffer from spatial-information loss and inadequate feature-representation issues. Spatial Information refers to information having location-based relation with other information (14).

Typically, after the training phase, during which neural network algorithms are provided with a large volume of relevant target data to hone their inference capabilities, and rewarded for correct responses in order to optimize performance, they’re essentially fixed. But Hasani’s team (21) developed a means by which his “liquid” neural net can adapt the parameters for “success” over time in response to new information, which means that if a neural net tasked with perception on a self-driving car goes from clear skies into heavy snow, for instance, it’s better able to deal with the shift in circumstances and maintain a high level of performance.

There are many decisions that could affect the results, by example, sometimes output could be affected due to the scaling method used because of the data distribution and whether standardization or normalization is chosen as the scaling method or maybe the data distribution is unknown and we don’t know what to choose, this is when using some metrics and visualizations previous to training could help us to choose the correct scaling method, when in doubt use the two scaling methods and compare the results with several of the metrics and benchmarking methods. But, in general, standardization is used when data is normally distributed whereas normalization technique does not assume any specific distribution.

There are many machine learning algorithms to choose from and in order to know whether the predictions for a given algorithm are good or not a well characterized dataset could be used but in lack of a well defined dataset a baseline prediction algorithm could help us establish a starting point for our datasets for comparison when evaluating other methods. A baseline is the lower limit defined by a set of benchmarking metrics, in the current form of *Bench-ML* the baseline prediction algorithm would be Decision Trees and the evaluated methods should be better than the baseline. The traditional baseline method “zero rule” would be way too basic and potentially confuse some of the predictions in undefined and complicated datasets. The question is, how much better than the baseline the metrics or predictions should be?. Using a variety of methods could help us identify limits or boundaries of how well the method or model is supposed to perform. Also, a variety of methods helps to know which methods work better and in which situations as sometimes a few methods will come at the top, but not always, and these surprises open the door to improvements in current or top of the line methods.

*Error rate* metric is implemented in Bench-ML and is monitored in order to keep it as low as possible; the two major sources of error are bias and variance. When these two are under control it is possible to build accurate models but in practice when bias is low - variance is high and vice versa because there is a trade-off between the two. If data is divided into training set and validation set there will be an error score for each and plotting the two error scores in learning curves is possible to see how the error changes as the training set size increases and the total error is a function of bias and variance.

**Image 1.**
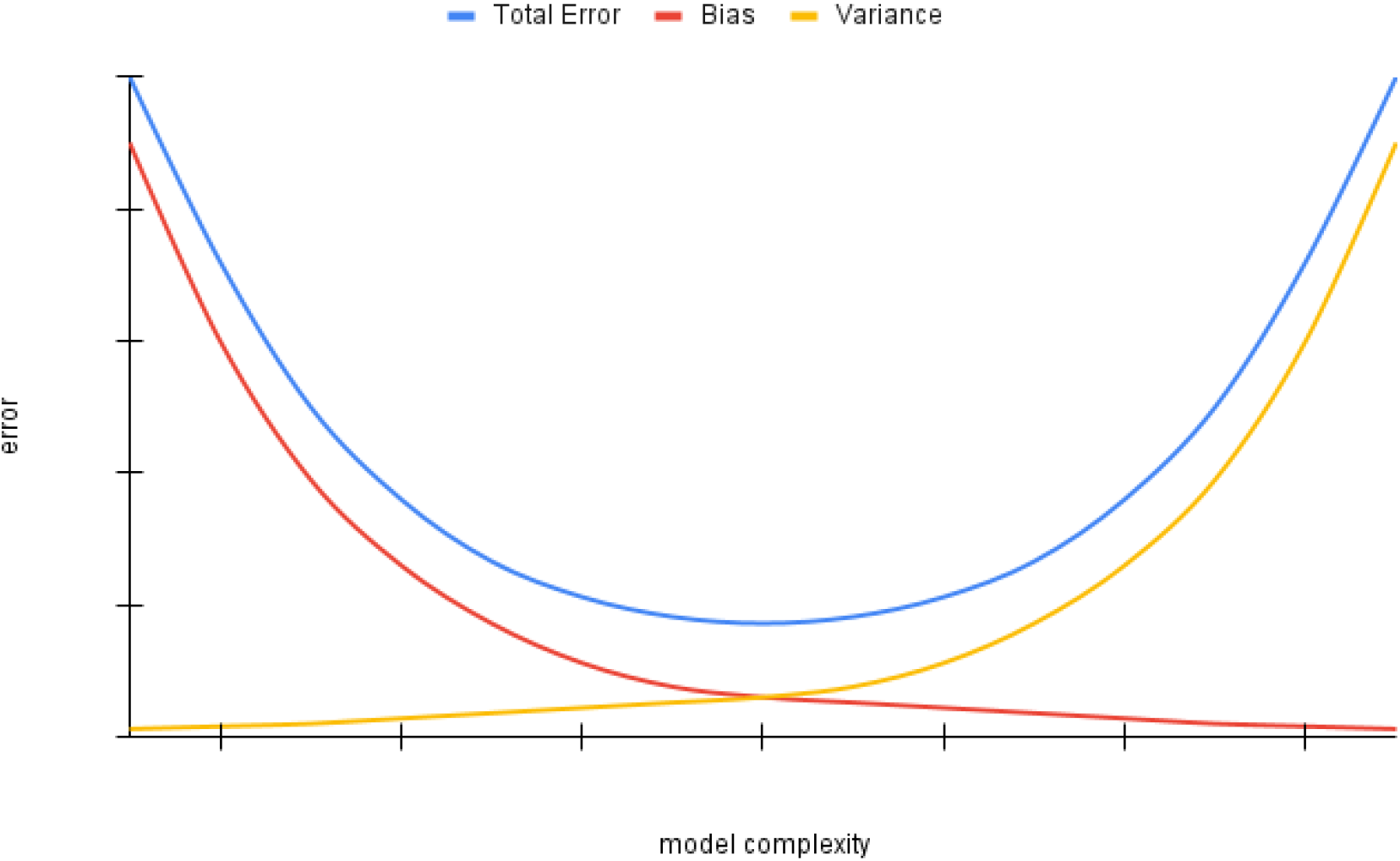
Total error is a function of Bias and Variance

**Image 2.**
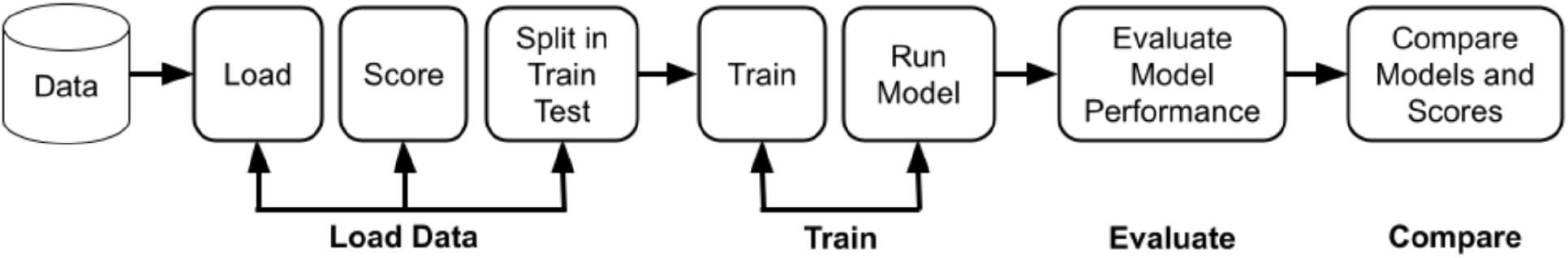
Bench-ML block design of the four major steps

**Image 3.**
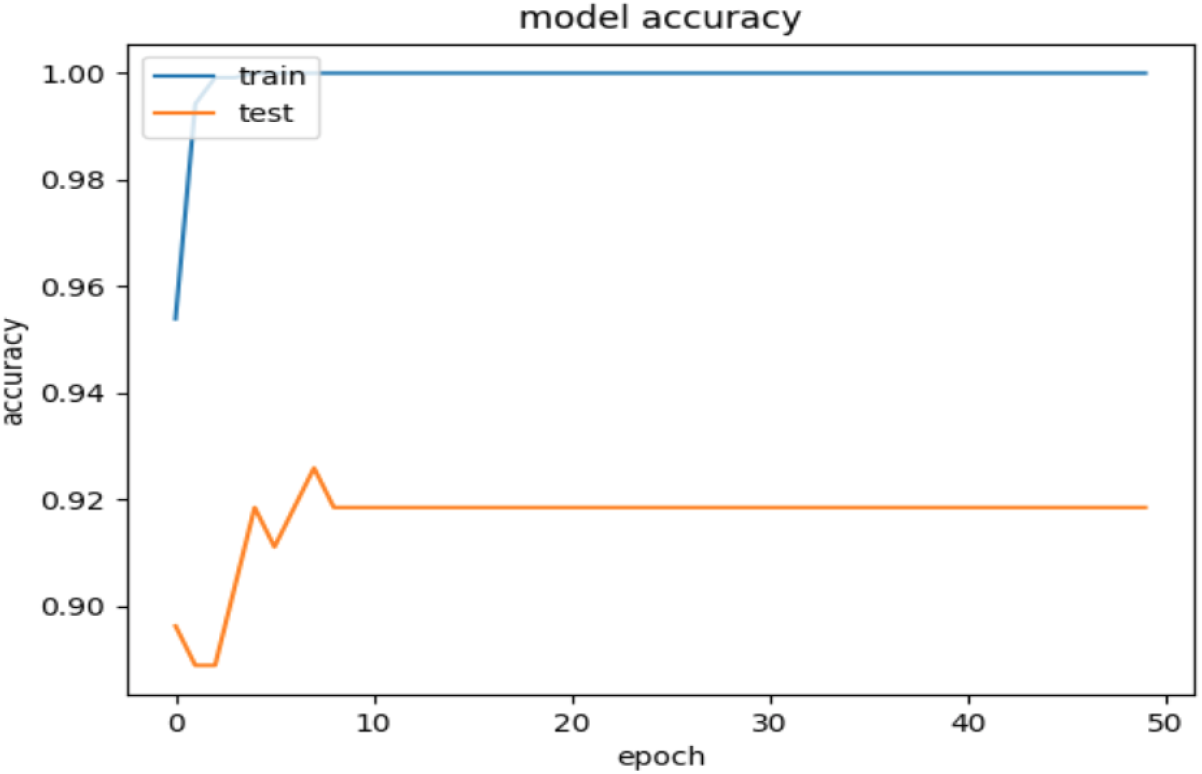
Model accuracy of GTEx of 6 tissue types using 100 features

**Image 4.**
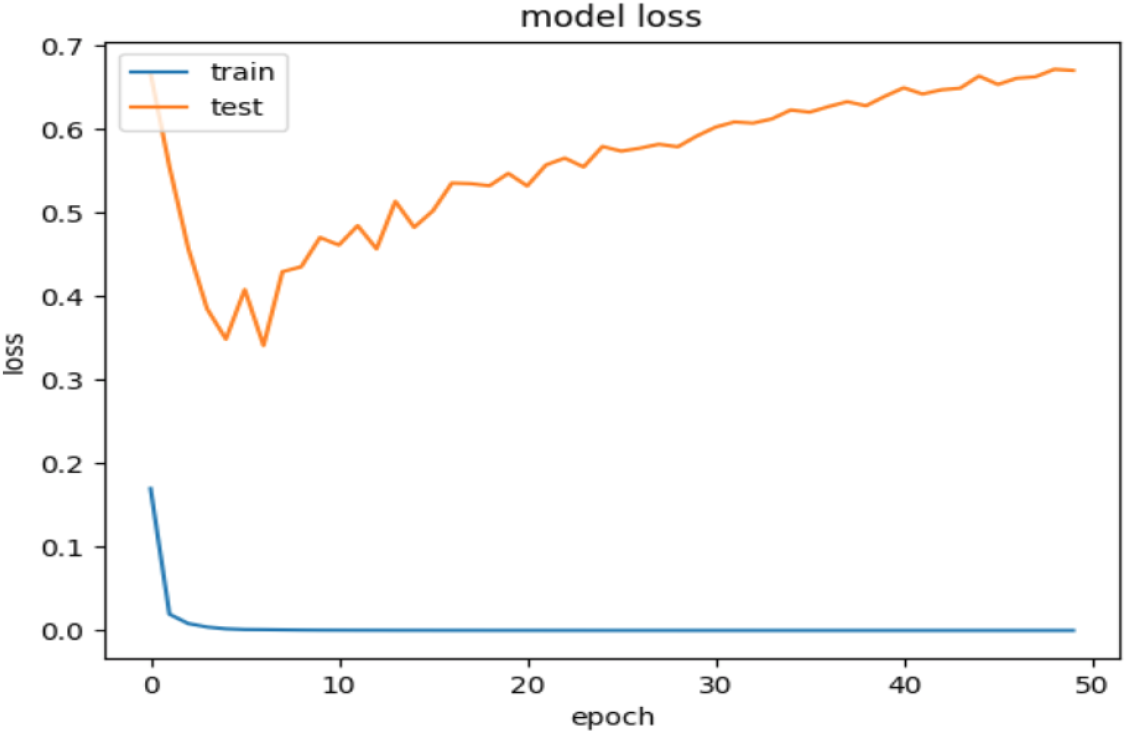
Model loss of GTEx of 6 tissue types using 100 features

**Image 5.**
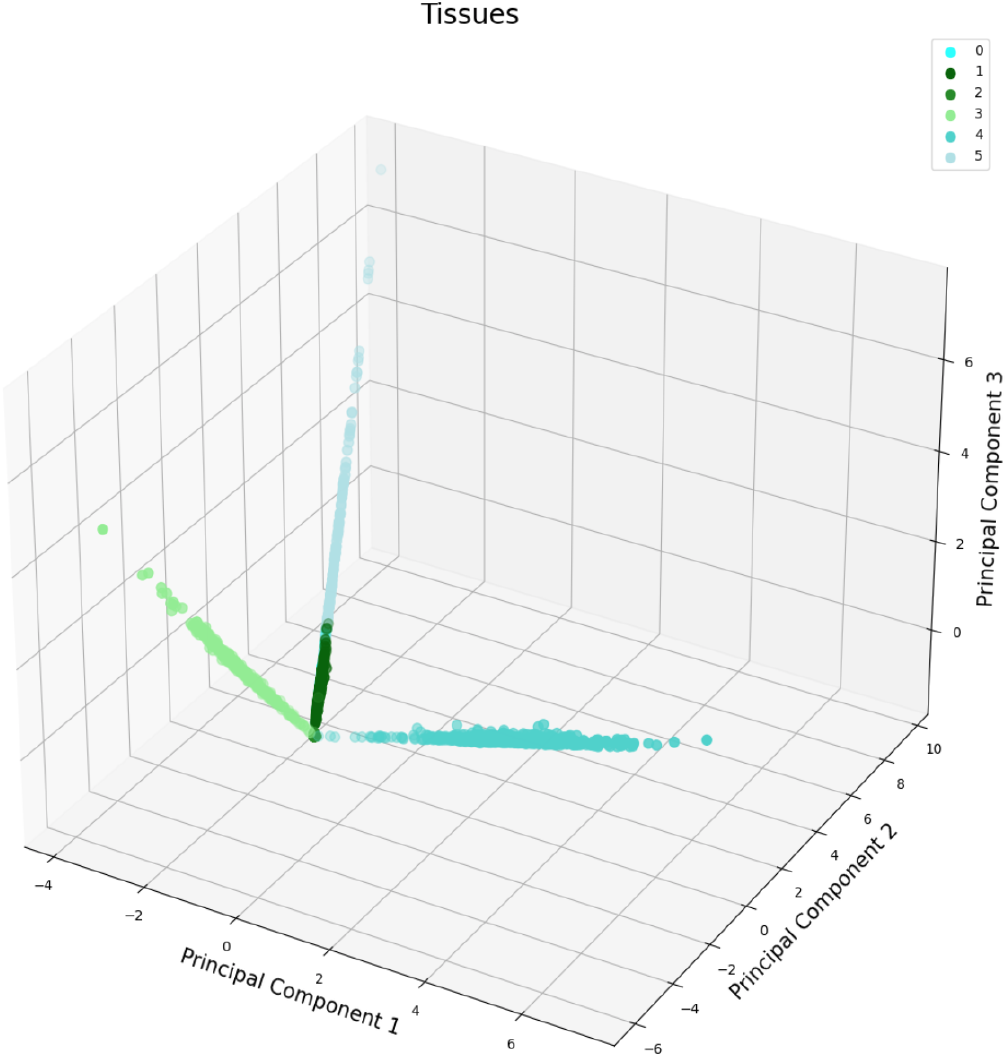
Example of Principal Component Analysis of GTEx with 6 tissue types using 100 features

**Image 6.**
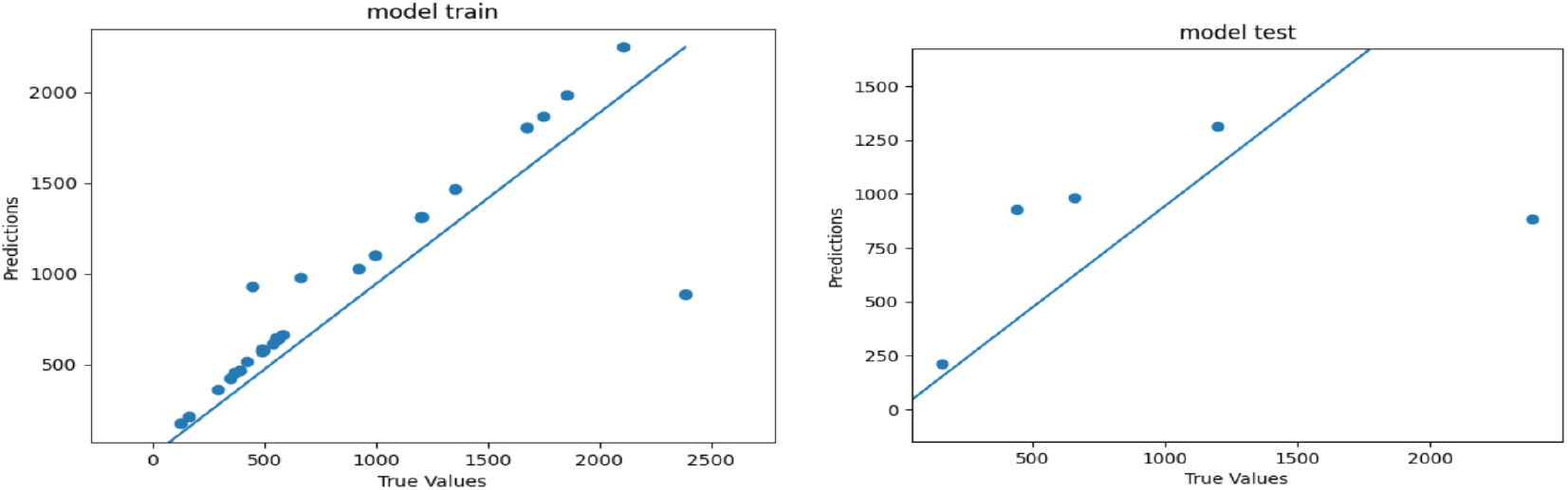
Example of regression error of the model with training and test data

**Image 7.**
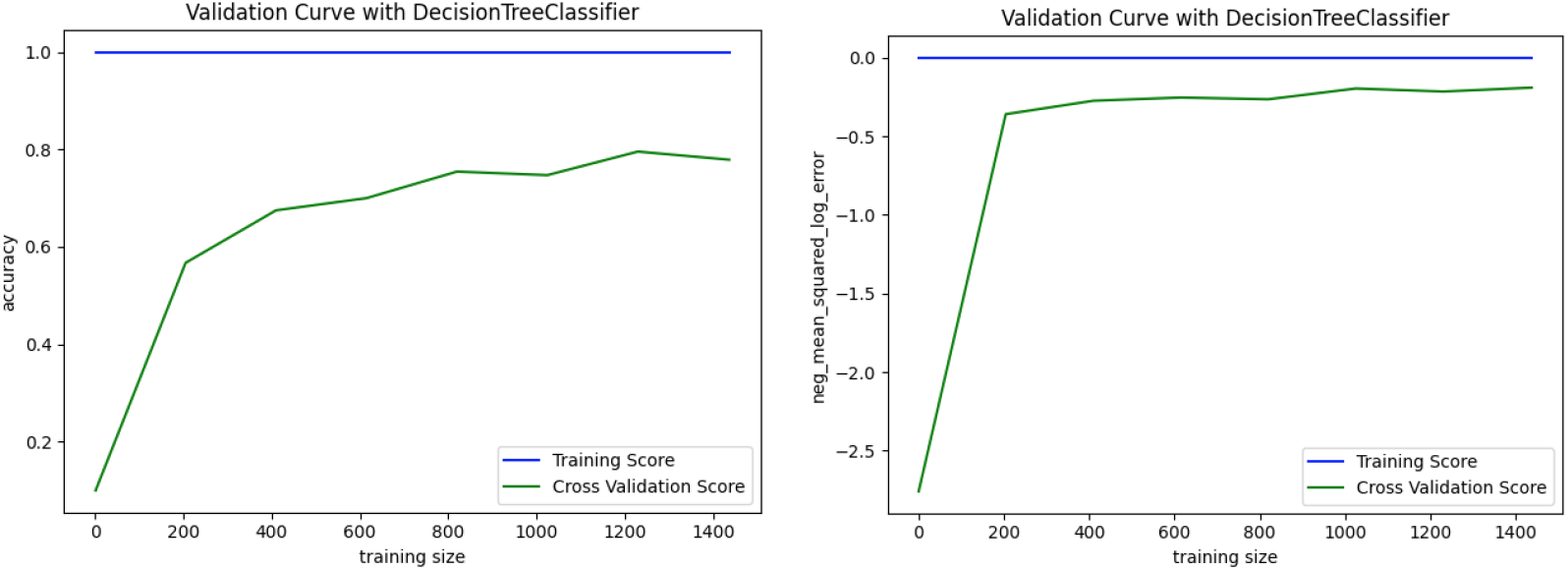
Accuracy and Model loss evaluated with Decision Trees

**Image 8.**
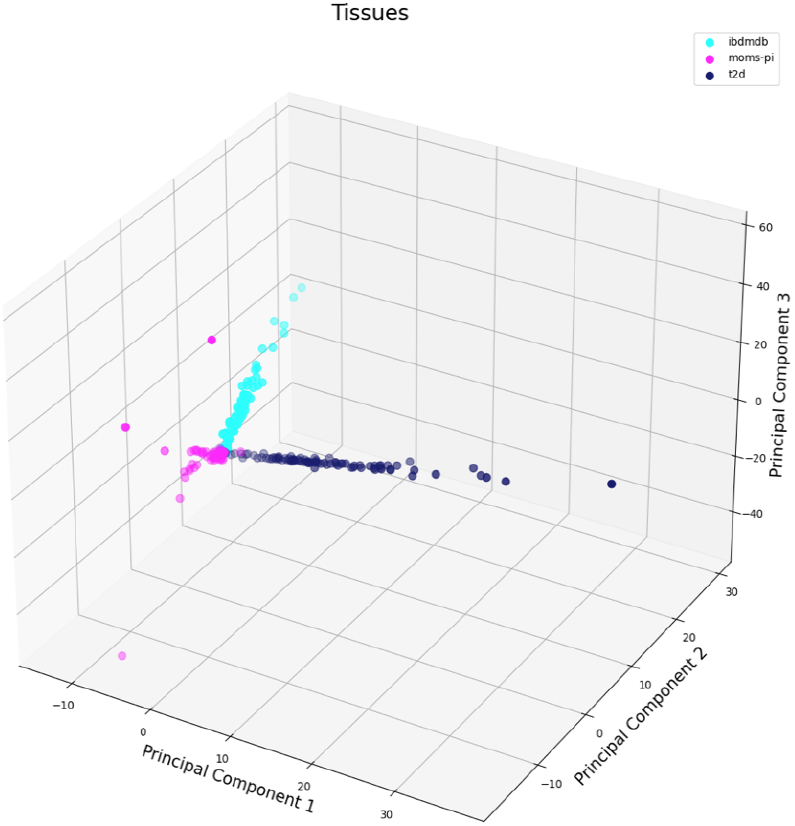
PCA for iHMP datasets to see how well samples cluster per disease type using all features

**Image 9.**
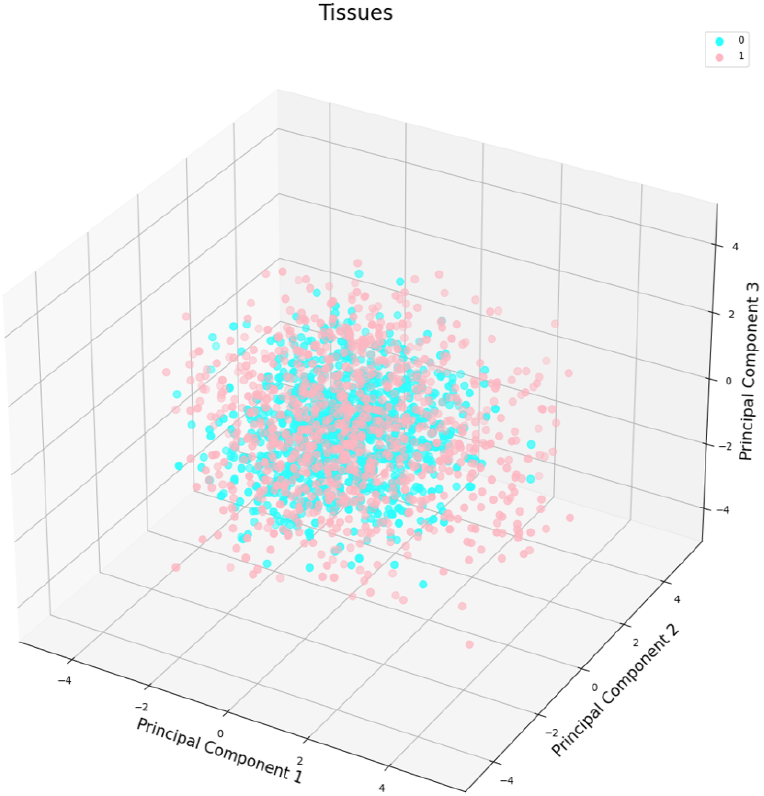
DNA Motifs PCA, it doesn’t show much separation between what is a motif and what is not.

**Image 10.**
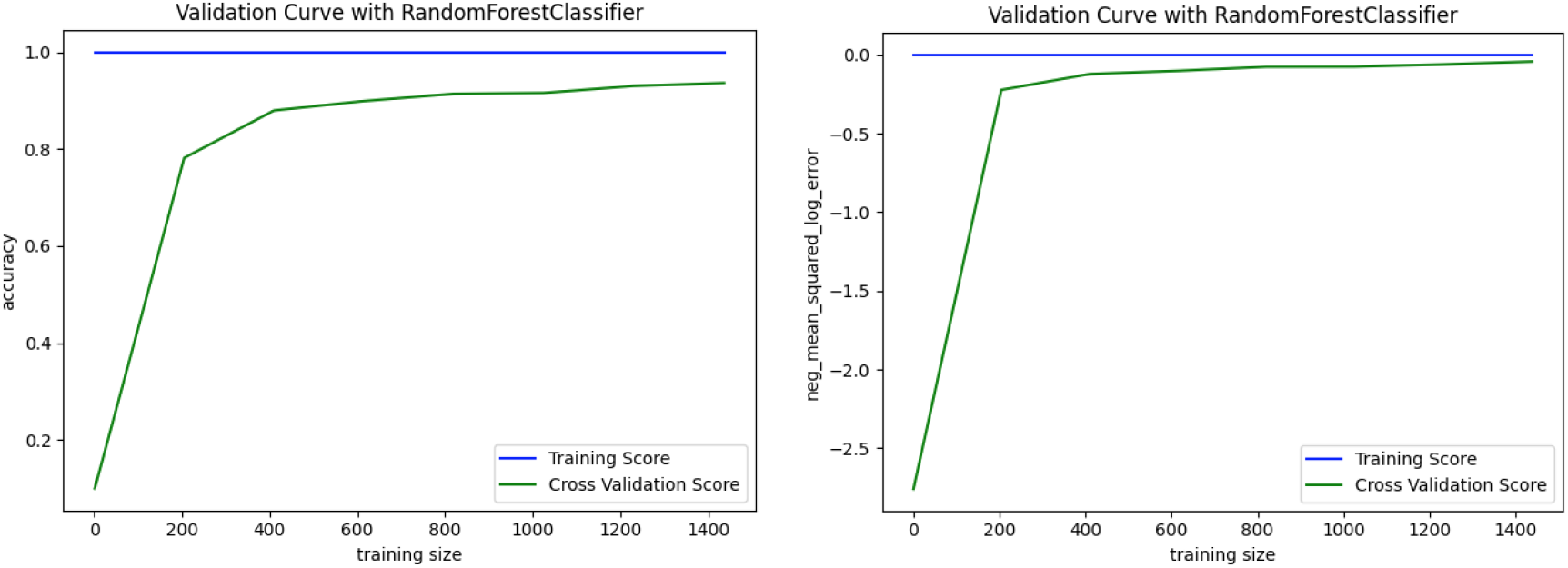
DNA Motifs accuracy and model loss with Random Forest trained using 100 features.

**Image 11.**
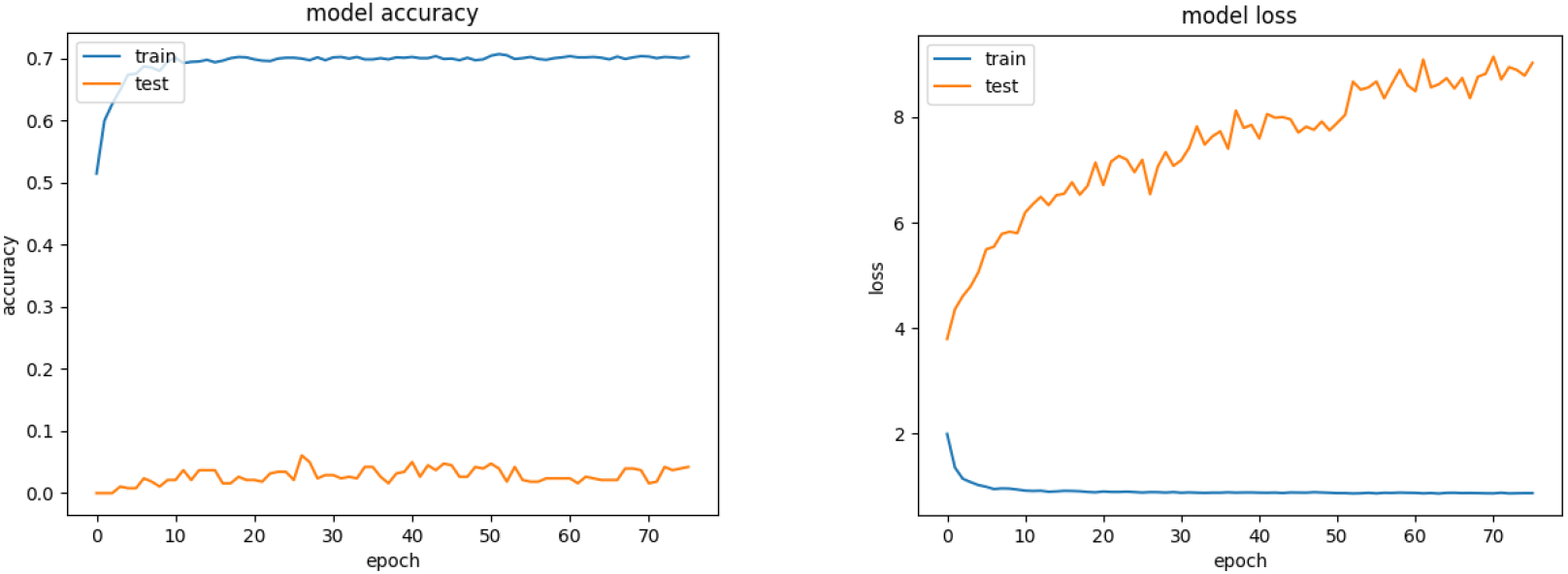
Accuracy and model loss for the TCGA Dataset

**Image 12.**
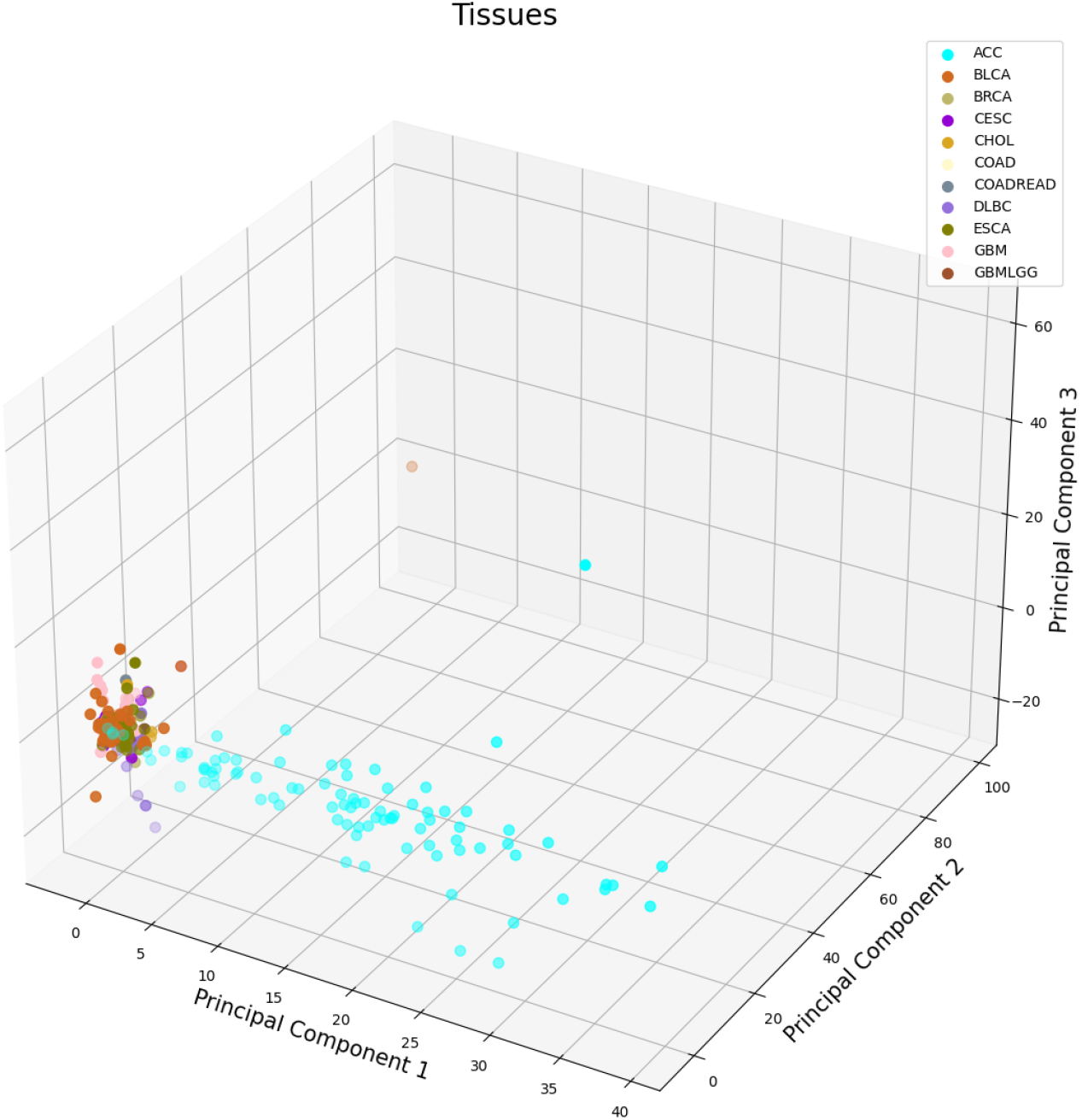
Principal Component Analysis for the TCGA Dataset

**Image 13.**
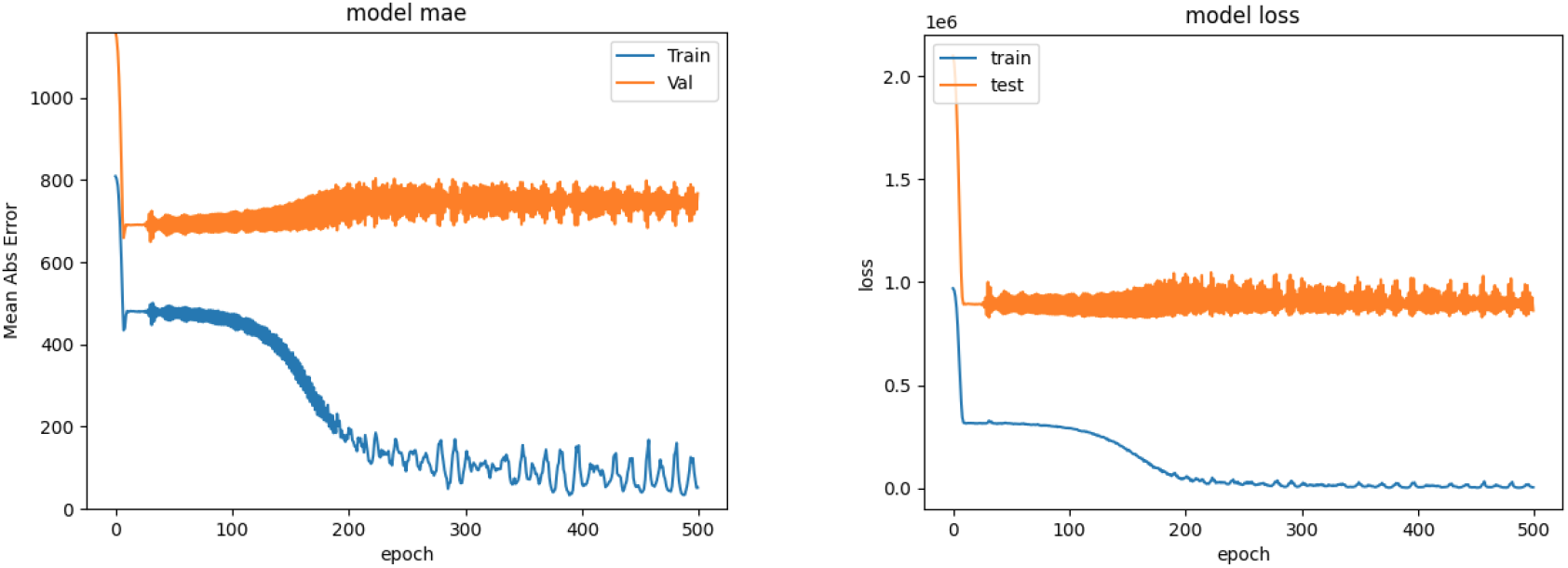
Accuracy and model loss for the TCGA Adrenocortical Carcinoma using Convolutional Neural Networks and all features

**Image 14.**
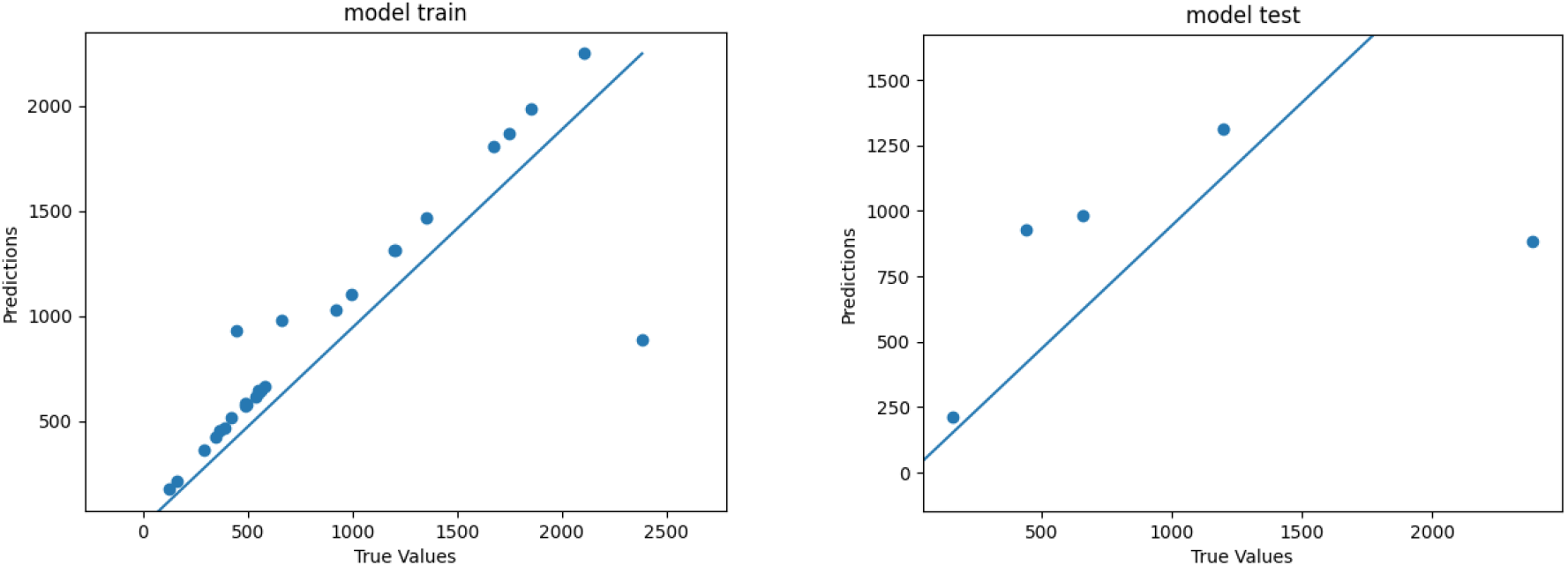
Regression for the TCGA Adrenocortical Carcinoma using Convolutional Neural Networks and all features

*Principal component analysis (PCA)* is also implemented in Bench-ML as a chart with two principal components (PC); PCA is a dimensionality reduction technique that arises from linear algebra and probability theory. It could use as many PCs as needed but it is traditionally used with two or three principal components; but when using three components it is recommended to use it in interactive mode.

## Design And Implementation

Bench-ML’s easy to use web interface allows users to select the dataset to be processed, define how to split the data into train and test, score features, select a machine learning method and its parameters so the user can easily see what the options are for each different method. It also has several options to display the model outputs in tables and visualize the results so the user can play with the parameters and Bench-ML can present a results matrix to compare each run. The user can learn from the predictions presented by Bench-ML and potentially generate new ideas on what could be done differently to improve the model or maybe learn that more data is needed for the method to perform at its best, or maybe the distribution is not what the model expected or something else given the variety of metrics, charts, models and methods available in Bench-ML.

Bench-ML could train and model different scenarios until it feels comfortable with the results; this recursive approach along with libraries to preprocess genomics data provides key benefits for testing and comparison, model building, feature scoring, reproducibility, visualization, use of standardized metrics and allows the user to play with the different methods, models and parameters until it finds the most generalizable and acceptable solution.

The complexity derived from dynamic and complicated genomics datasets might require analyzing hundreds of hyperparameters combinations and building many models for evaluation. Bench-ML uses a pipeline to execute analysis models to then evaluate, test and compare the many outcomes.

The lack of details of methods and algorithms undermines the scientific value of machine learning research, many machine learning methods are used as a black box and there are other obstacles that hinder transparent and reproducible AI research (23).

Bench-ML ensures reproducibility by recording all parameters and tools used, also, by freezing a copy of the train and test data to avoid variation that sometimes happens when randomly splitting datasets. Bench-ML machine learning analyses are completely reproducible with little or no variation on the end results. And reproducibility is important as it has become critical in machine learning research (23).

Bench-ML enables end-to-end machine learning analyses that begin with loading datasets and preprocessing primary biological data and ends with comparisons of machine learning models results that can make predictions using DNA sequences, variants, human gene expression or microorganisms expression of phenotypic attributes like cancer type, cancer survival progression, tissue expression type, and microbiome related diseases. End-to-end processing is possible because Bench-ML has many machine learning tools available for analyzing genomics, transcriptomics, metagenomics, metabolomics and many other types of biological data, as well as preprocessing biological data like converting DNA bases and categorical data to one hot encoding. It can also process any other type of data such as non-biological data as long as it conforms to a matrix of features and a label.

Bench-ML supports seven major steps in machine learning: loading and managing data, preprocessing, scoring, splitting, training, evaluation, and comparison of results by integrating several machine learning libraries, along with additional visualization and conversion tools including some custom ad hoc libraries built specifically for Bench-ML. Some of the most relevant libraries are TensorFlow which is an open source framework and a symbolic math library based on dataflow and differentiable programming for deep learning processing with a particular focus on training and inference of deep neural networks; Keras is a powerful and easy to use library and API for developing and evaluating deep learning models; Scikit-learn provides the foundation for most of the steps and machine learning libraries used within Bench-ML. Additional libraries are included to meet key needs for machine learning in genomics and biology including one hot encoding, feature scoring, feature selection, approaches for identifying imbalanced classes, and modeling approaches using Decision Trees, Random Forest, Support Vector Machines (SVM), and Deep Learning for prediction and regression of non-parametric data as well as Gaussian Naive Bayes for normally distributed datasets.

### DATA PREPROCESSING

In machine learning, data preprocessing is the first step towards improving the quality of data and the value of the information. The quality and value that can be derived from the data directly affects the ability of our model to learn and predict. The steps involved in data preprocessing are data loading, one-hot encoding, feature scoring, feature selection, balance classes, and split in train and test. Splitted datasets are saved as files so those could be used for model training to make reproducibility more consistent.

*There are two file types* used when loading data into Bench-ML:

● Comma separated values (CSV)
● Tab separated values (TSV)

*The data types* that can be processed in Bench-ML are:

**Table 1.** Genomics Data types

| Data Type | Description |
| --- | --- |
| DNA sequences | it takes bases in sequences as an input and converts them to one-hot encoding |
| DNA variants | it can take SNPs as an input and converts them to one-hot encoding |
| Gene Expression / Transcriptomics | it takes genes and a number that represents the gene being over or under express |
| Microbiome 16S, Metabolomics or Transcriptomics | it takes microorganisms id, names, or taxonomies and a number that represents whether the microorganism is over or under express or a quantity |
| Other non-biological data | it can also take any other type of data as long as it conforms to a matrix of features and a label |

**Table 2.** Example Datasets

| Name | Data Type | Description | URL |
| --- | --- | --- | --- |
| Genotype Tissue Expression (GTEx) | Gene Expression | Predicts the tissue type using gene expression data from 56 different tissue types | https://benchmarkmachinelearning.com/static/benchmark/images/gtex_6_100.tsv |
| Motifs Dataset | DNA Sequences | Predicts whether a DNA sequence data is a motif or not | https://benchmarkmachinelearning.com/static/benchmark/images/motifs.tsv |
| Iris Dataset | Other (non biological data) | Predicts the flower species from three different flower species | https://benchmarkmachinelearning.com/static/benchmark/images/iris.csv |

**Table 3.** Training Machine Learning Methods in Bench-ML

| Method name | Parametric | Deep Learning | Ensemble |
| --- | --- | --- | --- |
| *Decision Trees (DT)* | No | No | No |
| DT is a supervised learning method used for classification and regression. The goal is to create a model that predicts the value of a target variable by learning simple decision rules inferred from the data features. In Bench-ML this is considered the baseline method although sometimes it could perform better than some of the other methods |  |  |  |
| *Random Forest (RF)* | No | No | Yes |
| is an ensemble learning method for classification, regression and other tasks that combines many classifiers to provide solutions to complex problems. It operates by constructing a multitude of decision trees at training time. For classification tasks, the output of the random forest is the class selected by most trees and for regression tasks, the mean or average prediction of the individual trees is returned. Is suppose to perform better than decision trees because random forests tend to correct for overfitting to their training set and it generally outperforms decision trees |  |  |  |
| *Support Vector Machines (SVM)* | No | No | No |
| is a supervised learning linear model that can solve linear and non-linear problems with significant accuracy using less computation power compared to other methods; it can analyze data for classification and regression analysis. The algorithm creates lines or hyperplanes; these hyperplanes are decision boundaries in an N-dimensional space that distinctly classifies the data points into classes. The support vectors are data points that are closer to the hyperplane and influence the position and orientation of the hyperplane to maximize the margin of the classifier |  |  |  |
| *Gaussian Naive Bayes* | Yes | No | No |
| is for data that follows a normal distribution also known as gaussian distribution. It could perform better than other methods if the data is normally distributed where the probability distribution is symmetric about the mean and the data near the mean is more frequent in occurrence than the data far from the mean |  |  |  |
| *Fully Connected Neural Networks (FCNN)* | No | Yes | No |
| In this architecture all nodes or neurons in one layer are connected to the nodes in the next layer. It allows the selection of a model to use from three different models. |  |  |  |
| *Convolutional Neural Networks (CNN)* | No | Yes | No |
| are designed for processing structured arrays of data and automatically and adaptively learn spatial hierarchies of features through a backpropagation algorithm by using multiple building blocks, such as convolution layers, pooling layers, and fully connected layers. It allows the selection of a model to use from three different models. |  |  |  |

**Table 4.** GTEx Confusion Matrix with 6 tissue types using 100 features. Support refers to the number of samples used for testing/validation per class. Confusion matrix columns are samples 0 through 5

| Classification Report |  |  |  |  | Confusion Matrix |  |  |  |  |  |
| --- | --- | --- | --- | --- | --- | --- | --- | --- | --- | --- |
| CLASS | PRECISION | RECALL | F1-SCORE | SUPPORT |  |  |  |  |  |  |
| 0 | 0.6667 | 0.4000 | 0.5000 | 5 | **2** | 3 | 0 | 0 | 0 | 0 |
| 1 | 0.9091 | 0.9375 | 0.9231 | 32 | 1 | **30** | 0 | 0 | 0 | 1 |
| 2 | 1.0000 | 1.0000 | 1.0000 | 72 | 0 | 0 | **72** | 0 | 0 | 0 |
| 3 | 1.0000 | 1.0000 | 1.0000 | 73 | 0 | 0 | 0 | **73** | 0 | 0 |
| 4 | 1.0000 | 1.0000 | 1.0000 | 131 | 0 | 0 | 0 | 0 | **131** | 0 |
| 5 | 0.9667 | 1.0000 | 0.9831 | 29 | 0 | 0 | 0 | 0 | 0 | **29** |

**Table 5.** Example of comparison of methods and parameters for the GTEx dataset

| ID | MODEL ID | MODEL | SCORED | ACCURACY | EPOCHS | epochs | filters | kernels | poolsize | activation | activation_output | optimizer | loss | learning_rate | model_option | dropout | earlystop | DEL |
| --- | --- | --- | --- | --- | --- | --- | --- | --- | --- | --- | --- | --- | --- | --- | --- | --- | --- | --- |
| 236 | 217 | SVM | DT-all | 0.9912 |  |  |  |  |  |  |  |  |  |  |  |  |  |  |
| 237 | 217 | SVM | DT-10 | 0.9912 |  |  |  |  |  |  |  |  |  |  |  |  |  |  |
| 238 | 217 | SVM | DT-50 | 0.9912 |  |  |  |  |  |  |  |  |  |  |  |  |  |  |
| 239 | 217 | SVM | RF-all | 0.9912 |  |  |  |  |  |  |  |  |  |  |  |  |  |  |
| 240 | 217 | SVM | RF-10 | 0.9912 |  |  |  |  |  |  |  |  |  |  |  |  |  |  |
| 241 | 217 | SVM | RF-50 | 0.9912 |  |  |  |  |  |  |  |  |  |  |  |  |  |  |
| 258 | 220 | Conv1D | RF-10 | 0.9854 | 50 | 32 | 2 | 1 |  | relu | softmax | Adam | categorical_crossentropy | 0.003 | 1 | 0.20 | 25 |  |
| 255 | 220 | Conv1D | DT-10 | 0.9854 | 50 | 32 | 2 | 1 |  | relu | softmax | Adam | categorical_crossentropy | 0.003 | 1 | 0.20 | 25 |  |
| 254 | 220 | Conv1D | DT-all | 0.9854 | 50 | 32 | 2 | 1 |  | relu | softmax | Adam | categorical_crossentropy | 0.003 | 1 | 0.20 | 25 |  |
| 253 | 219 | FC | RF-50 | 0.9825 | 50 | 32 |  |  |  | relu | softmax | Adam | categorical_crossentropy | 0.003 | 2 | 0.20 | 25 |  |
| 251 | 219 | FC | RF-all | 0.9825 | 50 | 32 |  |  |  | relu | softmax | Adam | categorical_crossentropy | 0.003 | 2 | 0.20 | 25 |  |
| 257 | 220 | Conv1D | RF-all | 0.9825 | 50 | 32 | 2 | 1 |  | relu | softmax | Adam | categorical_crossentropy | 0.003 | 1 | 0.20 | 25 |  |
| 256 | 220 | Conv1D | DT-50 | 0.9795 | 50 | 32 | 2 | 1 |  | relu | softmax | Adam | categorical_crossentropy | 0.003 | 1 | 0.20 | 25 |  |
| 249 | 219 | FC | DT-10 | 0.9795 | 50 | 32 |  |  |  | relu | softmax | Adam | categorical_crossentropy | 0.003 | 2 | 0.20 | 25 |  |

**Table 6.**
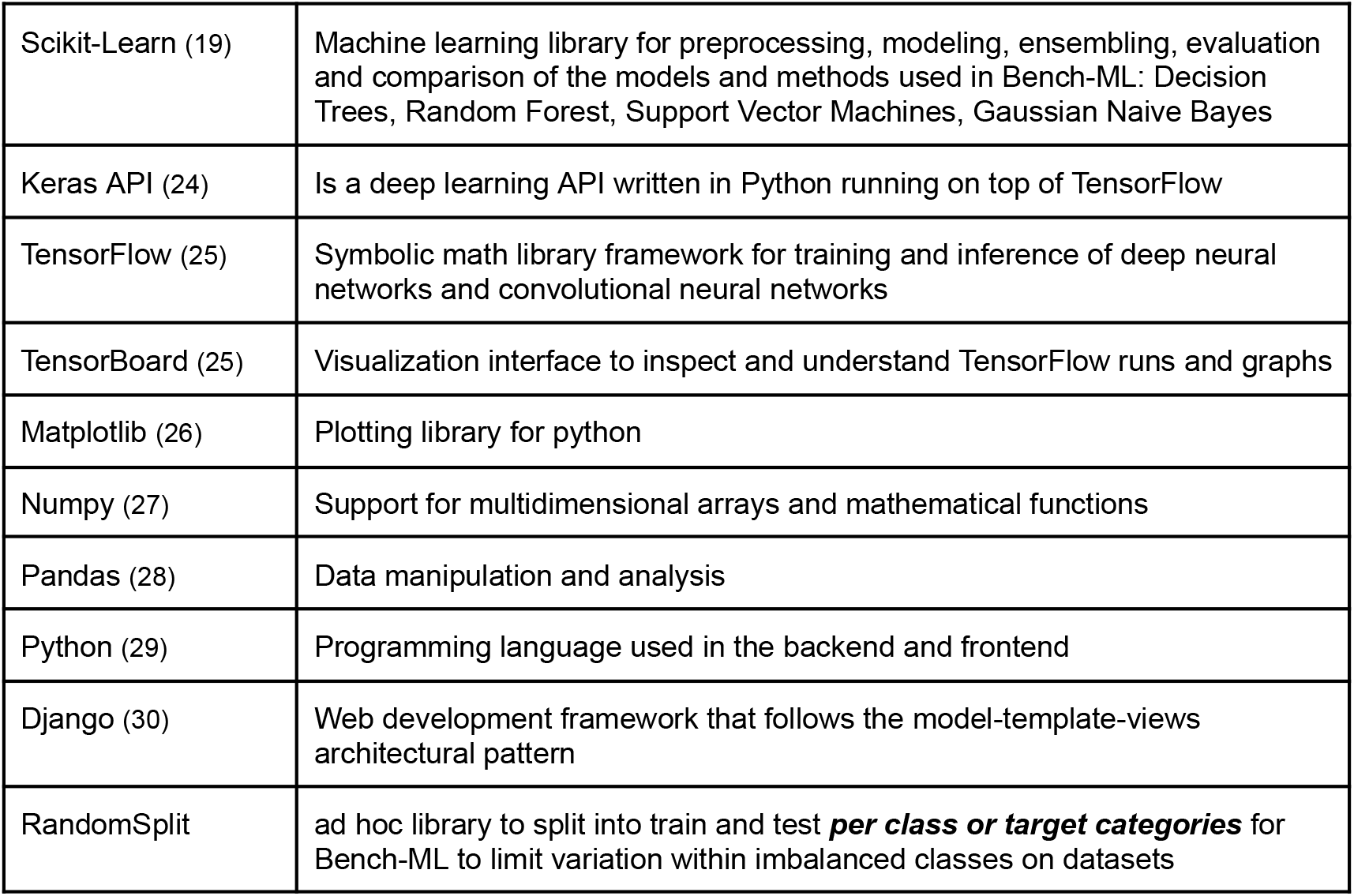
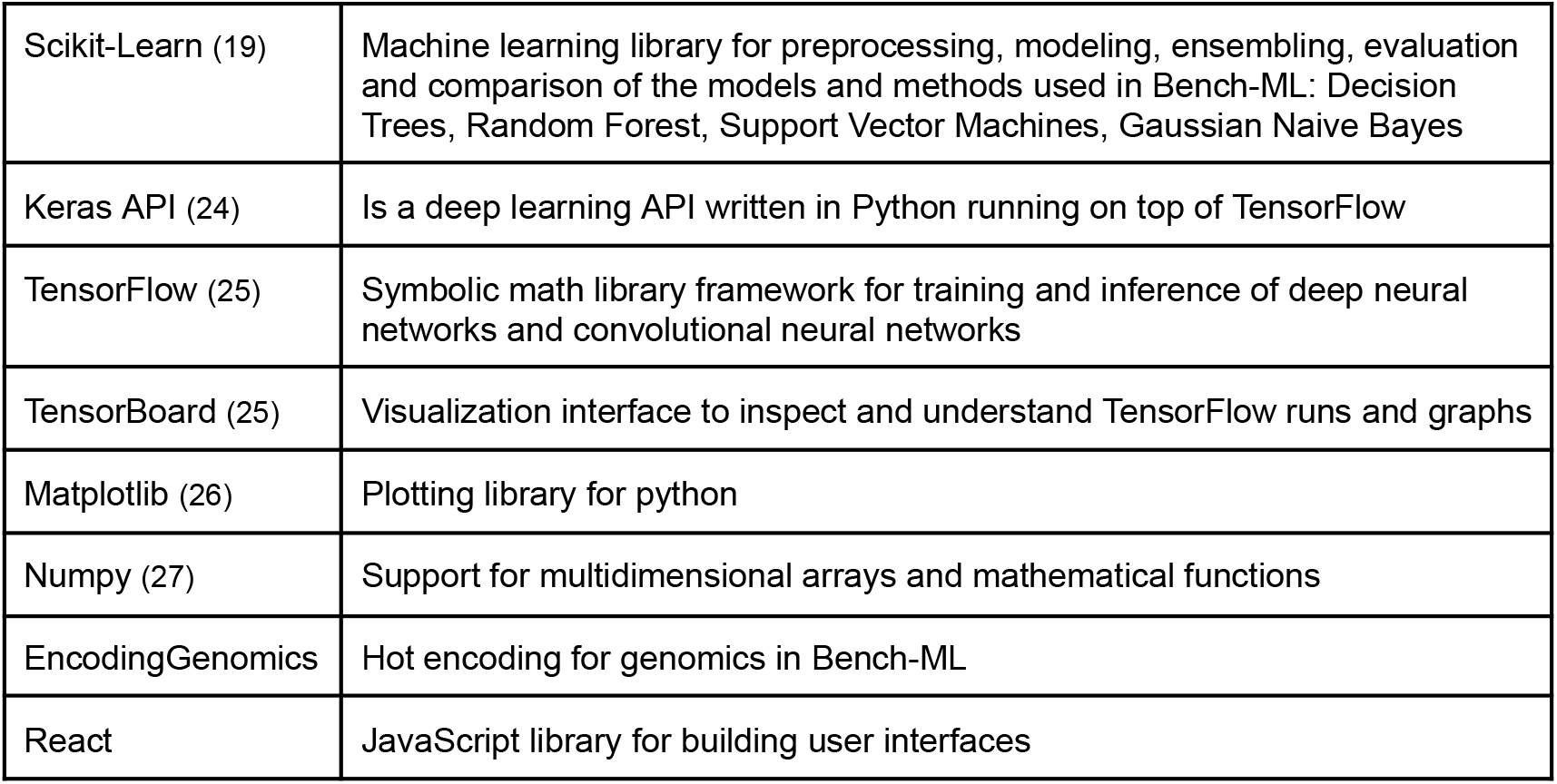
Software libraries integrated into Bench-ML

**Table 7.**
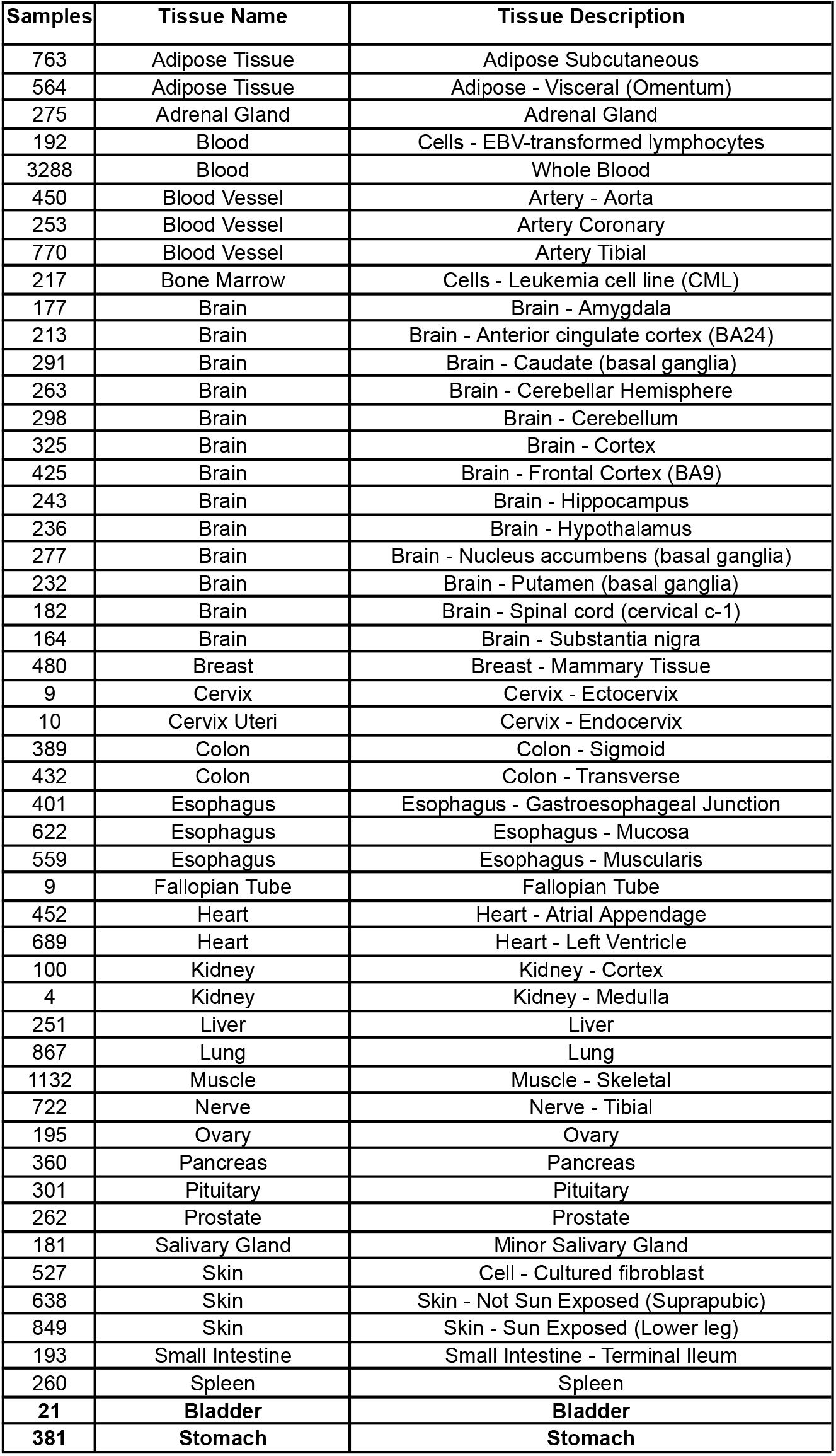
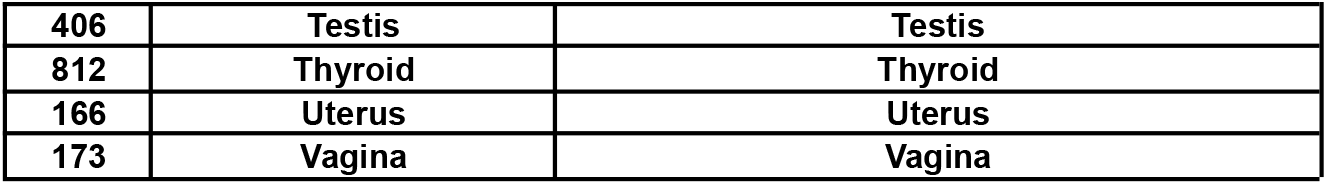
In the example dataset only 6 tissue types were used (in bold) out of 55 available

| Samples | Tissue Name | Tissue Description |
| --- | --- | --- |
| 763 | Adipose Tissue | Adipose Subcutaneous |
| 564 | Adipose Tissue | Adipose - Visceral (Omentum) |
| 275 | Adrenal Gland | Adrenal Gland |
| 192 | Blood | Cells - EBV-transformed lymphocytes |
| 3288 | Blood | Whole Blood |
| 450 | Blood Vessel | Artery - Aorta |
| 253 | Blood Vessel | Artery Coronary |
| 770 | Blood Vessel | Artery Tibial |
| 217 | Bone Marrow | Cells - Leukemia cell line (CML) |
| 177 | Brain | Brain - Amygdala |
| 213 | Brain | Brain - Anterior cingulate cortex (BA24) |
| 291 | Brain | Brain - Caudate (basal ganglia) |
| 263 | Brain | Brain - Cerebellar Hemisphere |
| 298 | Brain | Brain - Cerebellum |
| 325 | Brain | Brain - Cortex |
| 425 | Brain | Brain - Frontal Cortex (BA9) |
| 243 | Brain | Brain - Hippocampus |
| 236 | Brain | Brain - Hypothalamus |
| 277 | Brain | Brain - Nucleus accumbens (basal ganglia) |
| 232 | Brain | Brain - Putamen (basal ganglia) |
| 182 | Brain | Brain - Spinal cord (cervical c-1) |
| 164 | Brain | Brain - Substantia nigra |
| 480 | Breast | Breast - Mammary Tissue |
| 9 | Cervix | Cervix - Ectocervix |
| 10 | Cervix Uteri | Cervix - Endocervix |
| 389 | Colon | Colon - Sigmoid |
| 432 | Colon | Colon - Transverse |
| 401 | Esophagus | Esophagus - Gastroesophageal Junction |
| 622 | Esophagus | Esophagus - Mucosa |
| 559 | Esophagus | Esophagus - Muscularis |
| 9 | Fallopian Tube | Fallopian Tube |
| 452 | Heart | Heart - Atrial Appendage |
| 689 | Heart | Heart - Left Ventricle |
| 100 | Kidney | Kidney - Cortex |
| 4 | Kidney | Kidney - Medulla |
| 251 | Liver | Liver |
| 867 | Lung | Lung |
| 1132 | Muscle | Muscle - Skeletal |
| 722 | Nerve | Nerve - Tibial |
| 195 | Ovary | Ovary |
| 360 | Pancreas | Pancreas |
| 301 | Pituitary | Pituitary |
| 262 | Prostate | Prostate |
| 181 | Salivary Gland | Minor Salivary Gland |
| 527 | Skin | Cell - Cultured fibroblast |
| 638 | Skin | Skin - Not Sun Exposed (Suprapubic) |
| 849 | Skin | Skin - Sun Exposed (Lower leg) |
| 193 | Small Intestine | Small Intestine - Terminal Ileum |
| 260 | Spleen | Spleen |
| 21 | **Bladder** | **Bladder** |
| 381 | **Stomach** | **Stomach** |
| **406** | **Testis** | **Testis** |
| **812** | **Thyroid** | **Thyroid** |
| **166** | **Uterus** | **Uterus** |
| **173** | **Vagina** | **Vagina** |

**Table 8.**
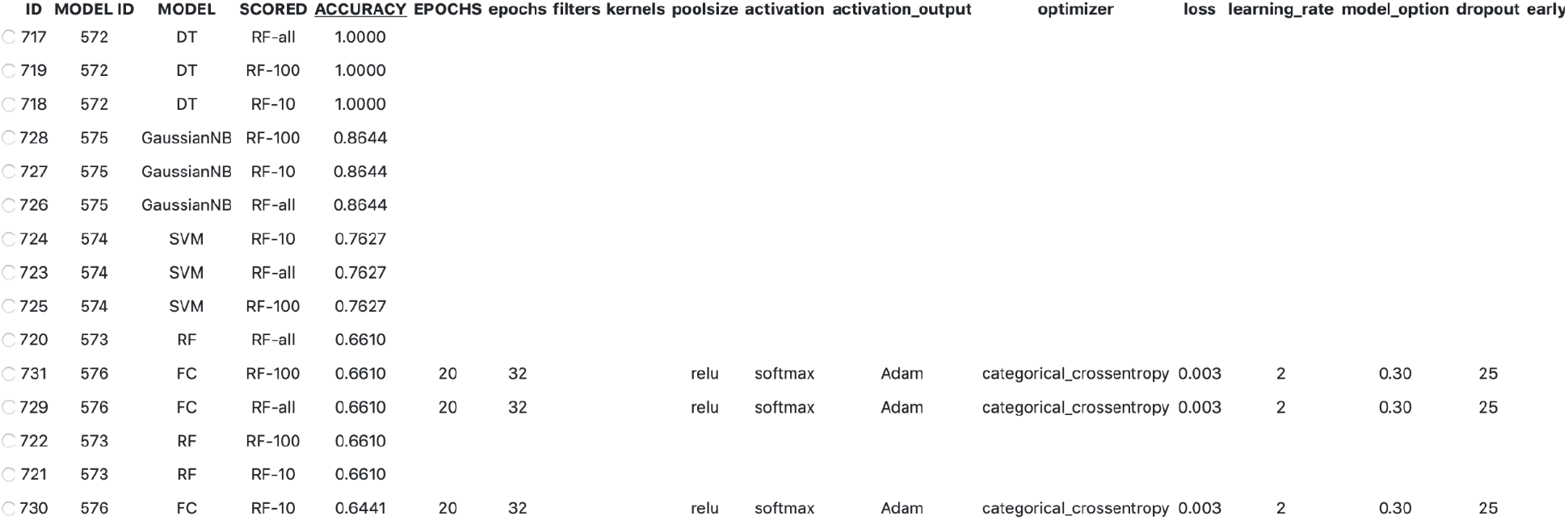
Comparison of methods results for iHMP

**Table 9.** iHMP type 2 diabetes (0), inflammatory bowel diseases (1), pregnancy and preterm birth (2) evaluated with Decision Trees

| Classification Report |  |  |  |  | Confusion Matrix |  |  |
| --- | --- | --- | --- | --- | --- | --- | --- |
| CLASS | PRECISION | RECALL | F1-SCORE | SUPPORT |  |  |  |
| 0 | 1.0000 | 1.0000 | 1.0000 | 19 | 19 | 0 | 0 |
| 1 | 1.0000 | 1.0000 | 1.0000 | 20 | 0 | 20 | 0 |
| 2 | 1.0000 | 1.0000 | 1.0000 | 20 | 0 | 0 | 20 |

**Table 10.** Predict Motifs Comparison of Methods

| ID | MODEL ID | MODEL | SCORED | ACCURACY | EPOCHS | epochs | filters | kernels | poolsize | activation | activation_output | optimizer | loss | learning_rate | model_option | dropout | earlys |
| --- | --- | --- | --- | --- | --- | --- | --- | --- | --- | --- | --- | --- | --- | --- | --- | --- | --- |
| 750 | 578 | RF | RF-100 | 0.8055 |  |  |  |  |  |  |  |  |  |  |  |  |  |
| 745 | 578 | RF | DT-100 | 0.8005 |  |  |  |  |  |  |  |  |  |  |  |  |  |
| 749 | 578 | RF | RF-50 | 0.7980 |  |  |  |  |  |  |  |  |  |  |  |  |  |
| 180 | 162 | SVM | RF-100 | 0.7980 |  |  |  |  |  |  |  |  |  |  |  |  |  |
| 175 | 162 | SVM | DT-100 | 0.7855 |  |  |  |  |  |  |  |  |  |  |  |  |  |
| 746 | 578 | RF | DT-all | 0.7781 |  |  |  |  |  |  |  |  |  |  |  |  |  |
| 751 | 578 | RF | RF-all | 0.7756 |  |  |  |  |  |  |  |  |  |  |  |  |  |
| 176 | 162 | SVM | DT-all | 0.7681 |  |  |  |  |  |  |  |  |  |  |  |  |  |
| 179 | 162 | SVM | RF-50 | 0.7681 |  |  |  |  |  |  |  |  |  |  |  |  |  |
| 744 | 578 | RF | DT-50 | 0.7681 |  |  |  |  |  |  |  |  |  |  |  |  |  |
| 174 | 162 | SVM | DT-50 | 0.7681 |  |  |  |  |  |  |  |  |  |  |  |  |  |
| 181 | 162 | SVM | RF-all | 0.7531 |  |  |  |  |  |  |  |  |  |  |  |  |  |
| 765 | 580 | Conv1D | DT-100 | 0.7406 | 20 | 32 | 2 | 1 | relu | softmax | Adam | categorical_crossentropy | 0.003 | 1 | 0.30 | 25 |  |
| 766 | 580 | Conv1D | DT-all | 0.7382 | 20 | 32 | 2 | 1 | relu | softmax | Adam | categorical_crossentropy | 0.003 | 1 | 0.30 | 25 |  |
| 770 | 580 | Conv1D | RF-100 | 0.7207 | 20 | 32 | 2 | 1 | relu | softmax | Adam | categorical_crossentropy | 0.003 | 1 | 0.30 | 25 |  |
| 760 | 579 | GaussianNB | RF-100 | 0.7182 |  |  |  |  |  |  |  |  |  |  |  |  |  |
| 740 | 577 | DT | RF-100 | 0.7082 |  |  |  |  |  |  |  |  |  |  |  |  |  |
| 764 | 580 | Conv1D | DT-50 | 0.7082 | 20 | 32 | 2 | 1 | relu | softmax | Adam | categorical_crossentropy | 0.003 | 1 | 0.30 | 25 |  |
| 771 | 580 | Conv1D | RF-all | 0.7057 | 20 | 32 | 2 | 1 | relu | softmax | Adam | categorical_crossentropy | 0.003 | 1 | 0.30 | 25 |  |
| 755 | 579 | GaussianNB | DT-100 | 0.7032 |  |  |  |  |  |  |  |  |  |  |  |  |  |
| 759 | 579 | GaussianNB | RF-50 | 0.7007 |  |  |  |  |  |  |  |  |  |  |  |  |  |
| 761 | 579 | GaussianNB | RF-all | 0.6983 |  |  |  |  |  |  |  |  |  |  |  |  |  |
| 769 | 580 | Conv1D | RF-50 | 0.6958 | 20 | 32 | 2 | 1 | relu | softmax | Adam | categorical_crossentropy | 0.003 | 1 | 0.30 | 25 |  |
| 170 | 161 | FC | RF-100 | 0.6883 |  | 32 |  |  | relu | softmax | Adam | categorical_crossentropy | 0.0003 | 1 | 0.30 | 10 |  |

**Table 11.** Comparison of Methods for Cancer types in the TCGA Dataset

| ID | MODEL ID | MODEL | SCORED | ACCURACY | EPOCHS | epochs | filters | kernels | poolsize | activation | activation_output | optimizer | loss | learning_rate | model_option | dropout | earlystop | DEI |
| --- | --- | --- | --- | --- | --- | --- | --- | --- | --- | --- | --- | --- | --- | --- | --- | --- | --- | --- |
| ○ 1146 | 704 | FC | RF-500 | 0.5084 | 500 | 128 |  |  |  | relu | softmax | RMSprop | categorical_crossentropy | 0.0003 | 2 | 0.10 | 75 | 🗑️ |
| ○ 1158 | 708 | FC | RF-500 | 0.5084 | 500 | 512 |  |  |  | relu | softmax | Adam | mean_squared_logarithmic_error | 0.0003 | 2 | 0.10 | 75 | 🗑️ |
| ○ 1077 | 686 | FC | RF-500 | 0.5084 | 500 | 256 |  |  |  | relu | softmax | Adam | categorical_crossentropy | 0.0003 | 2 | 0.01 | 0 | 🗑️ |
| ○ 1075 | 685 | FC | RF-500 | 0.5042 | 2000 | 128 |  |  |  | relu | softmax | Adam | categorical_crossentropy | 0.0003 | 2 | 0.10 | 0 | 🗑️ |
| ○ 1137 | 701 | FC | RF-500 | 0.5042 | 2000 | 128 |  |  |  | relu | softmax | Adam | categorical_crossentropy | 0.0003 | 2 | 0.10 | 75 | 🗑️ |
| ○ 1154 | 707 | FC | RF-500 | 0.5042 | 500 | 512 |  |  |  | relu | softmax | Adam | kullback_leibler_divergence | 0.0003 | 2 | 0.10 | 75 | 🗑️ |
| ○ 1066 | 682 | FC | RF-500 | 0.5042 | 500 | 64 |  |  |  | relu | softmax | Adam | categorical_crossentropy | 0.003 | 2 | 0.10 | 25 | 🗑️ |
| ○ 1062 | 681 | FC | RF-500 | 0.5021 | 500 | 32 |  |  |  | relu | softmax | Adam | categorical_crossentropy | 0.003 | 2 | 0.10 | 25 | 🗑️ |
| ○ 1168 | 712 | Conv1D | RF-500 | 0.5021 | 500 | 512 | 8 | 4 |  | relu | softmax | Adam | mean_squared_logarithmic_error | 0.003 | 2 | 0.05 | 75 | 🗑️ |
| ○ 1019 | 666 | FC | RF-500 | 0.5000 | 20 | 32 |  |  |  | relu | softmax | Adam | categorical_crossentropy | 0.003 | 1 | 0.30 | 25 | 🗑️ |
| ○ 1072 | 684 | FC | RF-500 | 0.5000 | 2000 | 128 |  |  |  | relu | softmax | Adam | categorical_crossentropy | 0.0003 | 2 | 0.10 | 25 | 🗑️ |
| ○ 1161 | 709 | FC | RF-500 | 0.5000 | 500 | 512 |  |  |  | relu | softmax | Adam | mean_squared_logarithmic_error | 0.0003 | 2 | 0.10 | 0 | 🗑️ |
| ○ 1149 | 705 | FC | RF-500 | 0.5000 | 1000 | 512 |  |  |  | relu | softmax | Adam | categorical_crossentropy | 0.0003 | 2 | 0.10 | 75 | 🗑️ |
| ○ 1140 | 702 | FC | RF-500 | 0.5000 | 500 | 128 |  |  |  | relu | softmax | Adam | categorical_crossentropy | 0.0003 | 3 | 0.10 | 75 | 🗑️ |
| ○ 1059 | 680 | Conv1D | RF-500 | 0.4979 | 500 | 128 | 2 | 1 |  | relu | softmax | Adam | categorical_crossentropy | 0.0003 | 2 | 0.10 | 25 | 🗑️ |
| ○ 1079 | 686 | FC | RF-all | 0.4979 | 500 | 256 |  |  |  | relu | softmax | Adam | categorical_crossentropy | 0.0003 | 2 | 0.01 | 0 | 🗑️ |
| ○ 1057 | 679 | Conv1D | RF-500 | 0.4958 | 500 | 64 | 2 | 1 |  | relu | softmax | Adam | categorical_crossentropy | 0.0003 | 2 | 0.10 | 25 | 🗑️ |
| ○ 1010 | 663 | RF | RF-500 | 0.4937 |  |  |  |  |  |  |  |  |  |  |  |  |  | 🗑️ |
| ○ 1148 | 704 | FC | RF-all | 0.4916 | 500 | 128 |  |  |  | relu | softmax | RMSprop | categorical_crossentropy | 0.0003 | 2 | 0.10 | 75 | 🗑️ |
| ○ 1012 | 663 | RF | RF-all | 0.4916 |  |  |  |  |  |  |  |  |  |  |  |  |  | 🗑️ |
| ○ 1150 | 705 | FC | RF-3000 | 0.4916 | 1000 | 512 |  |  |  | relu | softmax | Adam | categorical_crossentropy | 0.0003 | 2 | 0.10 | 75 | 🗑️ |
| ○ 1007 | 662 | DT | RF-500 | 0.4916 |  |  |  |  |  |  |  |  |  |  |  |  |  | 🗑️ |
| ○ 1020 | 666 | FC | RF-3000 | 0.4916 | 20 | 32 |  |  |  | relu | softmax | Adam | categorical_crossentropy | 0.003 | 1 | 0.30 | 25 | 🗑️ |
| ○ 1155 | 707 | FC | RF-3000 | 0.4916 | 500 | 512 |  |  |  | relu | softmax | Adam | kullback_leibler_divergence | 0.0003 | 2 | 0.10 | 75 | 🗑️ |
| ○ 1013 | 664 | SVM | RF-500 | 0.4895 |  |  |  |  |  |  |  |  |  |  |  |  |  | 🗑️ |

**Table 12.** Classification and Confusion Matrix for the TCGA Dataset

| Classification Report |  |  |  |  | Confusion Matrix |  |  |  |  |  |  |  |  |  |  |
| --- | --- | --- | --- | --- | --- | --- | --- | --- | --- | --- | --- | --- | --- | --- | --- |
| CLASS | PRECISION | RECALL | F1-SCORE | SUPPORT |  |  |  |  |  |  |  |  |  |  |  |
| 0 | 0.8750 | 0.7778 | 0.8235 | 18 | **14** | 2 | 0 | 2 | 0 | 0 | 0 | 0 | 0 | 0 | 0 |
| 1 | 0.3571 | 0.5769 | 0.4412 | 26 | 0 | **15** | 8 | 2 | 0 | 0 | 0 | 0 | 1 | 0 | 0 |
| 2 | 0.5146 | 0.9898 | 0.6771 | 196 | 0 | 0 | **194** | 2 | 0 | 0 | 0 | 0 | 0 | 0 | 0 |
| 3 | 0.3200 | 0.2051 | 0.2500 | 39 | 1 | 7 | 22 | **8** | 0 | 0 | 0 | 0 | 1 | 0 | 0 |
| 4 | 0.6667 | 0.2857 | 0.4000 | 7 | 0 | 1 | 0 | 1 | **2** | 0 | 0 | 1 | 2 | 0 | 0 |
| 5 | 0.0000 | 0.0000 | 0.0000 | 31 | 0 | 0 | 31 | 0 | 0 | **0** | 0 | 0 | 0 | 0 | 0 |
| 6 | 0.0000 | 0.0000 | 0.0000 | 45 | 0 | 0 | 45 | 0 | 0 | 0 | **0** | 0 | 0 | 0 | 0 |
| 7 | 0.8000 | 0.4000 | 0.5333 | 10 | 0 | 0 | 5 | 1 | 0 | 0 | 0 | **4** | 0 | 0 | 0 |
| 8 | 0.3750 | 0.0811 | 0.1333 | 37 | 1 | 6 | 22 | 4 | 1 | 0 | 0 | 0 | **3** | 0 | 0 |
| 9 | 0.0000 | 0.0000 | 0.0000 | 58 | 0 | 10 | 43 | 4 | 0 | 0 | 0 | 0 | 1 | **0** | 0 |
| 10 | 0.0000 | 0.0000 | 0.0000 | 9 | 0 | 1 | 7 | 1 | 0 | 0 | 0 | 0 | 0 | 0 | **0** |

**Table 13.** Accuracy and model loss for the Iris Flower Types Dataset

| ID | MODEL ID | MODEL | SCORED | ACCURACY | EPOCHS | epochs | filters | kernels | poolsize | activation | activation_output | optimizer | loss | learning_rate | model_option | dropout | early |
| --- | --- | --- | --- | --- | --- | --- | --- | --- | --- | --- | --- | --- | --- | --- | --- | --- | --- |
| 101 | 101 | RF | DT-all | 1.0000 |  |  |  |  |  |  |  |  |  |  |  |  |  |
| 204 | 185 | FC | DT-all | 1.0000 | 10 | 28 |  |  |  | relu | softmax | Adam | categorical_crossentropy | 0.03 | 2 | 0.10 | 21 |
| 102 | 102 | SVM | DT-all | 1.0000 |  |  |  |  |  |  |  |  |  |  |  |  |  |
| 103 | 103 | GaussianNB | DT-all | 1.0000 |  |  |  |  |  |  |  |  |  |  |  |  |  |
| 105 | 105 | FC | DT-all | 1.0000 |  | 32 |  |  |  | relu | softmax | Adam | categorical_crossentropy | 0.003 | 2 | 0 | 25 |
| 107 | 107 | Conv1D | DT-all | 1.0000 |  | 32 | 2 | 1 |  | relu | softmax | Adam | categorical_crossentropy | 0.003 | 1 | 0 | 25 |
| 184 | 165 | Conv1D | DT-all | 1.0000 |  | 32 | 2 | 1 |  | relu | softmax | Adam | categorical_crossentropy | 0.003 | 1 | 0.30 | 25 |
| 97 | 97 | DT | DT-all | 0.9667 |  |  |  |  |  |  |  |  |  |  |  |  |  |
| 186 | 167 | Conv1D | DT-all | 0.9667 |  | 32 | 2 | 1 |  | relu | softmax | Adam | categorical_crossentropy | 0.003 | 1 | 35 | 25 |
| 190 | 171 | Conv1D | DT-all | 0.9667 |  | 32 | 2 | 1 |  | relu | softmax | Adam | categorical_crossentropy | 0.003 | 1 | 0.30 | 25 |
| 203 | 184 | Conv1D | DT-all | 0.9667 | 20 | 32 | 2 | 1 |  | relu | softmax | Adam | categorical_crossentropy | 0.003 | 1 | 0.30 | 25 |
| 205 | 186 | Conv1D | DT-all | 0.9667 | 21 | 21 | 2 | 1 |  | relu | softmax | Adam | categorical_crossentropy | 0.03 | 1 | 0.21 | 21 |
| 224 | 205 | Conv1D | DT-all | 0.9667 | 28 | 28 | 2 | 1 |  | relu | softmax | Adam | categorical_crossentropy | 0.0028 | 1 | 0.28 | 28 |
| 225 | 206 | Conv1D | DT-all | 0.9667 | 29 | 29 | 2 | 1 |  | relu | softmax | Adam | categorical_crossentropy | 0.0029 | 1 | 0.29 | 29 |
| 260 | 221 | Conv1D | DT-all | 0.9333 | 10 | 32 | 2 | 1 |  | relu | softmax | Adam | categorical_crossentropy | 0.003 | 1 | 0.20 | 25 |
| 189 | 170 | Conv1D | DT-all | 0.9000 |  | 32 | 2 | 3 |  | relu | softmax | Adam | categorical_crossentropy | 0.003 | 1 | 0.30 | 25 |
| 520 | 375 | FC | DT-all | 0.6333 | 20 | 32 |  |  |  | relu | softmax | Adam | categorical_crossentropy | 0.003 | 2 | 0.30 | 25 |
| 209 | 190 | FC | DT-all | 0.6000 | 20 | 20 |  |  |  | relu | softmax | Adam | categorical_crossentropy | 0.0003 | 2 | 0.30 | 25 |

*Feature scoring*, many times the data is trained and evaluated with all the features or with a subset of the most important features for testing purposes. Bench-ML web interface has an option to define how many different combinations of features to use including using all features. I.e. The interface could take “all” features or a combination of a number of features like “all,10,50,100” features. For “all,10,50,100” the model will be run with “all”, with “10”, with “50”, and with “100” features. There are two methods for feature scoring and one could use one of the methods or both and the models will be run with all the combinations of features and methods for scoring; the methods for scoring are:

● Decision trees (DT), Bench-ML uses DT to score the features and to select only the most significant features.
● Random forests (RF), Bench-ML can also use RF for feature scoring in case a subset of the most important features is needed.

*Datasets could be split in train and test* according to the percent defined in Bench-ML, by default it uses 80% of the data for training and 20% for testing. The resulting files are saved so results will be more consistent when using different methods because it uses the same datasets.

There is an option that can be selected to balance the classes or targets or labels according to the training percent. I.e. 80% of class A will be used for training, 80% of class B, 80% of class C, etc.

*The regression flag* is used for datasets that need to calculate the measure of the relation between the mean value of one variable and corresponding values of other variables instead of using classification. For example, the TCGA cancer survival dataset uses regression in its model to predict how many days a person could survive a specific cancer type.

*Example datasets*, there are three example datasets available from the Bench-ML web interface that could be used to test the web application from data already prepared and available for download at https://benchmark machine learning.com/score/. These working datasets are examples of how data is expected to be feeded into Bench-ML:

### MODEL TRAINING

Bench-ML has six machine learning methods to choose from along with its corresponding hyperparameters; deep learning also has several predefined models to choose from. There is one ensemble method as well as non-parametric and parametric methods that expect data that is normally distributed, also, a deep learning method with fully connected nodes and a deep learning method better suited for data in structured arrays. Within Bench-ML, Decision Trees could be used as the baseline method as it is the simplest solution from all the methods available.

### EVALUATE METHODS AND PARAMETERS

Bench-ML helps evaluate input files, feature scoring methods, models and training methods along with hyperparameters chosen using a variety of metrics, charts and matrices; It allows comparison of feature selection, feature scoring methods, training models and methods and parameters for an input file.

Bench-ML benchmarks each class or label; it provides a confusion matrix with metrics like precision, recall, f1-score and number of samples used to evaluate each label performance. In machine learning it is common to provide a single accuracy score but in genomics a single label (class or disease) not performing could make a difference for a group of people, so it is important to learn about those deficiencies and find prediction alternatives for those classes like in the example below where Class 0 has a low precision and recall.

The training and validation data is shown separately in the accuracy charts. If model accuracy for the test data stays the same after some epochs it might need more data or a different model in order to improve but if the accuracy keeps improving it might need to increase the number of epochs as it could go higher. In the image below it hints that it needs more data to improve accuracy for both data sets.

The training and validation data is also shown separately in the loss charts. Model loss for the test data will usually increase because after some epochs it doesn’t generalize anymore but it could give us an idea in which epoch to stop using the Bench-ML “early stop” parameter.

Bench-ML uses principal component analysis to identify groups of samples, it might identify which groups overlap or whether a group could be divided into more than one or if the data is easily divided or it isn’t very easy to identify groups.

For regression Bench-ML also uses two charts to display the model with training data as well as with test data to visualize how well the test data aligns compared to the training data.

### COMPARE METHODS AND PARAMETERS

Bench-ML allows easy comparisons of feature scoring methods, training methods and hyperparameters in one table. It orders the results by accuracy score with the most accurate method and parameters on the top to quickly visualize the best performing combinations for the dataset being analyzed and identify methods and parameters that haven’t been tested and could potentially improve performance.

### DOCUMENTATION AND SOFTWARE USED

Example datasets are available at https://benchmarkmachinelearning.com/score

Documentation is available at:

● For loading and preparing data https://benchmarkmachinelearning.com/docscore
● For training https://benchmarkmachinelearning.com/doctrain

Bench-ML code and tool repositories are available at https://github.com/rlopeztec/bench-ml

## Results and Discussion

The utility of Bench-ML was demonstrated in five use cases: Predict tissue type from the GTEx dataset, Predict microbiome phenotype using the Human Microbiome dataset, Discover DNA binding motifs from an example dataset, Cancer survival regression from the TCGA dataset, and Predict flower types from an example dataset; the flower types dataset was only used to demonstrate that Bench-ML can process non-biological data.

The “Compare Methods and Parameters” section from above provides links to complete analyses and results so that all analyses can be fully reproduced on the Bench-ML web interface. All analyses were performed on the public Bench-ML web server at https://benchmarkmachinelearning.com. All workflows, data and results can be accessed via a web browser and analyses can be reproduced directly, although some analyses will take a long time to reproduce.

### Predicting Tissue Type (GTEx)

The Genotype Tissue Expression (GTEx) program established a data resource and tissue bank to study the relationship between genetic variants (inherited changes in the DNA), regulation and gene expression (how genes are turned on and off) in multiple tissue types and across individuals. GTEx consists of 55 non-diseased tissue types across nearly 1,000 individuals, primarily for molecular assays including Whole Genome Sequencing (WGS), Whole Exome Sequencing (WES), and RNA-Seq to evaluate regulation and gene expression (39).

Bench-ML was used to predict from six different tissue types from the GTEx RNA-Seq dataset consisting of 56,201 samples to illustrate the utility from Bench-ML. Six tissue types were selected with very different amounts of samples to evaluate if the varying number of training data has an effect on prediction accuracy; the six tissue types are marked in blue in the table below.

One model was applied to the GTEx dataset using 100 features and 6 tissue types across the following machine learning methods: SVM, DT, RF, GNB, CNN, and DNN.

### Predicting Microbiome Phenotype (From iHMP Dataset)

The NIH Human Microbiome Project (HMP) has been carried out over ten years and two phases to provide resources, methods, and discoveries that link interactions between humans and their microbiomes to health-related outcomes. The first phase (HMP1) was launched in 2007 and the second phase or integrative Human Microbiome Project (iHMP) was launched in 2014 (31).

Bench-ML used the data from the iHMP dataset that included the following phenotypes: Pregnancy and Preterm birth, Inflammatory Bowel Disease (IBD), and Type 2 Diabetes (T2D).

### Discover DNA Binding Motifs (Example Dataset)

Bench-ML can discover DNA binding motifs. A DNA motif is a short sequence with a recurrent pattern in DNA that is presumed to have a biological function, often indicating sequence specific binding sites for proteins (8).

### Cancer type classification (from the TCGA dataset)

The Cancer Genome Atlas Program (TCGA) is a joint effort between NCI and NHGRI that molecularly characterized over 20,000 samples consisting of primary cancer and matched normal samples spanning 33 cancer types (40).

Bench-ML can predict cancer types from TCGA real patients using six different machine learning methods and compare the different methods to see what performs better.

### Cancer survival regression (from the TCGA dataset)

Bench-ML predicted the number of days a cancer patient could survive from The Cancer Genome Atlas Program (TCGA) using regression from Deep Learning (42).

### Iris Flower Types (Non-biological example)

The iris flower is a non-biological dataset which is frequently used in machine learning.

### Monitor ML Models

Monitoring is a well defined practice for many software systems although for machine learning is not as easy because an ML system needs to retrain and rerun the model with the new dataset and compare it with previous results to understand how accurate the predictions are over time. Bench-ML helps monitor this by maintaining a copy of the model, versions, and parameters.

Because of the data distribution the ML system could require changes on the model structure. To test these models a well characterized dataset is partitioned into training and test data then the models are tuned until it gets to the desired sensitivity and specificity. Continuous monitoring of changes could identify datasets that do not longer work as expected on certain models or it could highlight the need for different models for different reasons due to natural drifting that happens if the data is changing when (17):

● The relationship between the model and the input data changes
● The distribution of the data changes such that the model is less representative
● Changes in measurements or user base which changes the underlying meaning of variables
● A significant deviation in the distribution of the new set of predictions might indicate degradation in performance
● If the prediction probabilities are low this could indicate the model is struggling

Monitoring input data into the model to help track the deterioration of predictions (17) and student’s t-test or nonparametric Kolmogorov-Smirnov methods could be used to compare distributions:

● The distribution of features might deviate significantly from the training data over time
● The number of categories or classes of a model’s variable might increase over time

### Reproducibility

All of the analyses in the Bench-ML web tool are highly reproducible because it saves the data used for training and testing intact for later use as well as the corresponding parameters and versions used in the web application. When loading the data Bench-ML splits training and test data on a given percentage (default is 20%) these datasets are stored on the file system to be used in the same or other analysis with the same data for reproducibility purposes, this way it removes the randomness that occurs when splitting the data in test and training since randomness could modify the results at training or when evaluating the predictions with the test data. Bench-ML also stores and records all parameter settings and versions used on each analysis as well as the corresponding results.

The data splitted in training and test could be downloaded and used along with the parameters in the framework tensorflow to reproduce the same training model and results using the model structure selected in the web application as well as the parameters and versions of the software packages.

When preprocessing genomics datasets features could be in the thousands or millions of features. Bench-ML can preprocess the data to do feature scoring, it could use all features or a subset of the features. It scores the features using either decision trees of random forest then selects the features with the greatest scoring, this set of features are then stored in the file system so the files could be used to reproduce the analyses with minimum variability as oppose to re-score the features and select a different set of features given that decision trees and random forest could score features differently given that could pick a different feature when the features has the same importance and this process is stochastic or random in nature.

### Extensibility

Bench-ML already has many custom packages and software libraries but it can be extended, basically it can accept any number of custom methods and machine learning software packages and libraries. Bench-ML has a set of integrated tools from existing machine learning software packages and libraries as well as custom developed software packages for data preprocessing as well as libraries specific to genomics like one hot encoding de entire nucleotides sequences of DNA or variants like single nucleotide polymorphisms i.e. adenine (A), thymine (T), cytosine (C), and guanine (G). It also does feature scoring in case the user wants to have a reduced set of features and not all of them because in genomics the features could be thousands or even millions of features depending on how the features were defined. Bench-ML could also take care of the model definition, training, and evaluation of the model selected.

Bench-ML provides integration points where additional custom made software packages and machine learning libraries can be added. For example if a new machine learning library becomes available it could be added and tools such as model fit, model predict, and can be augmented so users can use models from this machine learning library. In previous sections custom modules were implemented for feature scoring using decision trees and random forest, data splitting in training and test, preprocessing, modeling, prediction, and evaluation.

### Availability and future directions

Often effects of hyperparameters on models are known but a better approach is to search different values and choose a subset that achieves the best performance. Bench-ML could Automate model’s evaluation and propose efficiency improvements by using automated hyper-parameter tuning or hyper parameter optimization (HPO), GridSearch defines a search

space as a grid of hyperparameter values and evaluates every position, RandomSearch which defines the search space as a bounded domain and randomly sample points in that domain, Bayesian Optimization directs a search of a global optimization building a probabilistic model of the objective function and Evolutionary Optimization which is a generic population-based metaheuristic optimization algorithm inspired by biological evolution.

Some models could benefit from algorithms to automatically detect unusual data points that differ significantly from other data points and potentially treat them differently. Some of those algorithms are: Elliptic Envelope which is machine learning based, Minimum Covariance Determinant uses statistical methods if the data has a gaussian distribution, IQR uses statistics, Isolation Forest that is a Tree based Algorithm effective in high dimensional data, and One-class SVM and Local Outlier Factor (LOF) that uses unsupervised machine learning algorithms.

*Learning architectures* or algorithms focus on training the model to make fewer predictions errors, it usually requires less data and converges faster. Neural Architecture Search (NAS) is a learning architecture and there are other techniques such as distillation used to improve the accuracy of a smaller model by learning to mimic a larger model like the Data Efficient Image Transformer (DeiT) developed by Facebook, it is a vision transformer built on a transformer specific knowledge distillation procedure to reduce training data requirements. Distillation allows neural networks to learn amongst themselves where a distillation token is a learned vector that flows through the network along with the transformed image data to significantly enhance the image classification performance with less training data.

K-fold cross validation is a resampling procedure commonly used to compare and select machine learning models using statistical methods on limited data and it uses a single parameter K to split the data in a number of groups.

## Conclusions

Bench-ML is a benchmarking and evaluation tool with custom made computational programs as well as existing software libraries with popular machine and deep learning methods that were used to properly check and validate models and parameters in genomics datasets where machine and deep learning are a very active development field. It also increases the reproducibility of the ML methods by providing a tool with the methods and all the parameters along with the dataset used for training and testing.

Four complex genomics datasets were used in Bench-ML by non machine learning developers and bioinformaticians to demonstrate how to preprocess genomics data into a format understandable by machine learning. It helped train and verify the performance and prediction accuracy of the models using a combination of parameters and methods on thousands of features and samples. Bench-ML also provided insights into the potential bias and limitations of the models or the data using tables to facilitate the comparison of models trained and executed to easily identify the best performing methods, models and parameters.

Bench-ML can process any dataset as long as it conforms to a matrix of features and a column of labels or classes and build models to validate their performance in non-biological datasets as demonstrated by using the Iris flowers dataset.

## Glossary

● Deep Learning, Neural networks with multiple layers of artificial neurons. The output of one layer is fed as input into the next layer to achieve greater flexibility.
● Back-propagation, A common method to train neural networks by updating its parameters (i.e., weights) by using the derivative of the network’s performance with respect to the parameters
● Feed-forward neural network, The most flexible class of neural networks, wherein each neuron can have arbitrary weights
● Convolutional neural network, A class of neural networks in which groups of artificial neurons are scanned across the input to identify translation-invariant patterns.
● Recurrent neural network, A class of neural networks with cycles that can process inputs of varying lengths.
● Autoencoder, A class of neural networks that performs nonlinear dimensionality reduction.
● in-sample data, is the data used to fit the model
● out-of-sample data, is data that wasn’t part of the model fitting and is used to forecast the performance

## Notes

https://benchmarkmachinelearning.com/

### Competing Interest Statement

The authors have declared no competing interest.

